# In vivo Safety and Immunoactivity of Oncolytic Jurona Virus in Hepatocellular Carcinoma: A Comprehensive Proteogenomic Analysis

**DOI:** 10.1101/2022.09.09.507330

**Authors:** Yuguo Zhang, Mulu Tesfay, Khandoker U. Ferdous, Mika Taylor, Musa Gabere, Camila C. Simoes, Chelsae Dumbauld, Oumar Barro, Alicia L. Graham, Charity L. Washam, Duah Alkam, Allen Gies, Jean Christopher Chamcheu, Stephanie D. Byrum, Steven R. Post, Thomas Kelly, Mitesh J. Borad, Martin J. Cannon, Alexei Basnakian, Bolni M. Nagalo

## Abstract

Oncolytic viruses can effectively unwrap a multimodal anti-tumor activity, encompassing a selective tumor cell killing and promoting a systemic anti-tumor immunity, making them a formidable foe against cancer. Among these, several members of the Rhabdoviridae family are particularly attractive as oncolytic agents due to their natural tumor selectivity and non-pathogenicity in humans. In this study, we demonstrated that intratumorally (IT) administration of Jurona virus (JURV), a novel oncolytic Rhabdovirus, induces dynamic tumor regression in human HCC xenograft and syngeneic models. Our data shows that IT injections of JURV trigger the recruitment and activation of cytotoxic T (CTLs) and decrease the tumor-associated macrophages (TAM) infiltration leading to tumor growth delay in both local and distant murine HCC tumors in a syngeneic model. Moreover, when administered concomitantly, JURV and anti-PD-1 therapy profoundly modulate the tumor microenvironment (TME) via enhanced infiltration of CTLs, suggesting that immune checkpoint blockade therapy could potentiate the immunomodulatory effect of JURV and potentially provide durable anti-tumor immunity. Our analysis of the molecular and cellular mechanism of JURV-medicated anti-cancer activity unveiled that JURV and anti-PD-1 antibodies activate different effectors of the immune system but have complementary anti-tumor activities. Furthermore, our results indicate that the abscopal effect induced by JURV is likely mediated by the mechanism regulating the T helper cell responses. Our work supports the further development of JURV as a novel immunovirotherapy platform for hepatocellular carcinoma.

## INTRODUCTION

Hepatocellular carcinoma (HCC) is a major cause of cancer related morbidity and mortality worlwide.^1^ Most HCC patients are diagnosed at advanced stages and are left with limited therapeutic options. Current approaches for advanced disease include cytotoxic therapies, targeted therapies, and immune checkpoint inhibitors. ^23^ Unfortunately, most patients with HCC do not achieve long-term disease control, making HCC one of cancer with the highest unmet clinical need globally.

Oncolytic viruses (OVs) are formidable foe against cancer; they replicate preferentially in tumor cells that are deficient in their anti-viral innate immune responses.^6, 7^ Given the multifaceted anti-cancer activities of OVs, which comprise direct tumor cell killing capabilities and immunomodulatory properties, many viruses, including members of *Rhabdoviridae*^2, 3^ family are becoming increasingly appealing in immuno-oncology.^1, 2^ A recent clinical study published in blood advances^3^ brought to light the long-awaited outcome of vesicular stomatitis virus (VSV), a vesiculovirus of the family Rhabdoviridae, encoding the interferon-beta transgene (VSV-IFNβ) for patients with advanced lymphoma, highlighting once again the incredible oncolytic potential of these viruses.

In this study, we reported that Jurona virus (JURV) ^4, 5^ a non-VSV vesiculovirus JURV can effectively induce oncolysis *in vitro,* in animal models of HCC and prime systemic anti-tumor immunity resulting in robust tumor regression in syngeneic and xenograft hepatocellular carcinoma (HCC) models. Co-administration of JURV with immune checkpoint blockade antibodies profoundly modulated the tumor microenvironment via increase infiltration of cytotoxic T cells. This timely and compelling study will provide ground for continued evaluation of the extraordinary oncolytic potential of JURV in first-in-human studies.

## RESULTS

### JURV potently infects and lyse human and murine HCC cell lines

Analysis of the complete genome of a laboratory attenuated JURV clone showed that the 10,993 bp genome displays typical *Rhabdoviridae* organization and consist of five major genes from the 3’ to 5’ antigenomic direction: nucleoprotein (JURV-N), phosphoprotein (JURV-P), matrix (JURV-M), glycoprotein (JURV-G), polymerase (JURV-L) (Supplemental Fig. 1a-c).^6^ The tumor-cell adapted JURV clone, wild-type VSV, and wild-type morreton virus (MORV) were used to infect monolayers of human and murine HCC cells at different multiplicity of infection (MOI) of 10, 1 and 0.1 (Fig. 1a-e). An MTS cell viability assay assessment of cytotoxic at 72 hours post-infection showed that susceptibility to viral killing and the virus-induced cell death was similar at MOI of 10 and 1 for these three viruses. However, we found a slight difference in virus-induced cell death in RILWT at an MOI of 0.1 between JURV, MORV, and VSV. To consolidate our findings, HCC cells were stained with crystal violet to assess the comparative cytotoxic effect of JURV or mock-infected HCC cells. We found that while HuH7 had residual (<50%) live adherent cells remaining after infection, HEP3B, PLC, HEPA 1-6, and RILWT (>90%) had a complete loss of adherent cells, suggesting that MTS may have slightly underestimated the oncolysis effect of JURV (Fig. 1f). Furthermore, analysis of JURV kinetics showed an exponential (∼ 1000-fold) increase in viral titers around 10 hours post-infection (Fig. 1g), indicating a high permissivity and potent replication capability of JURV in tumor cells. These results also highlight the importance of combining at least two methods for *in vitro* assessment of cell viability upon infection with an oncolytic virus.

**Figure 1.**
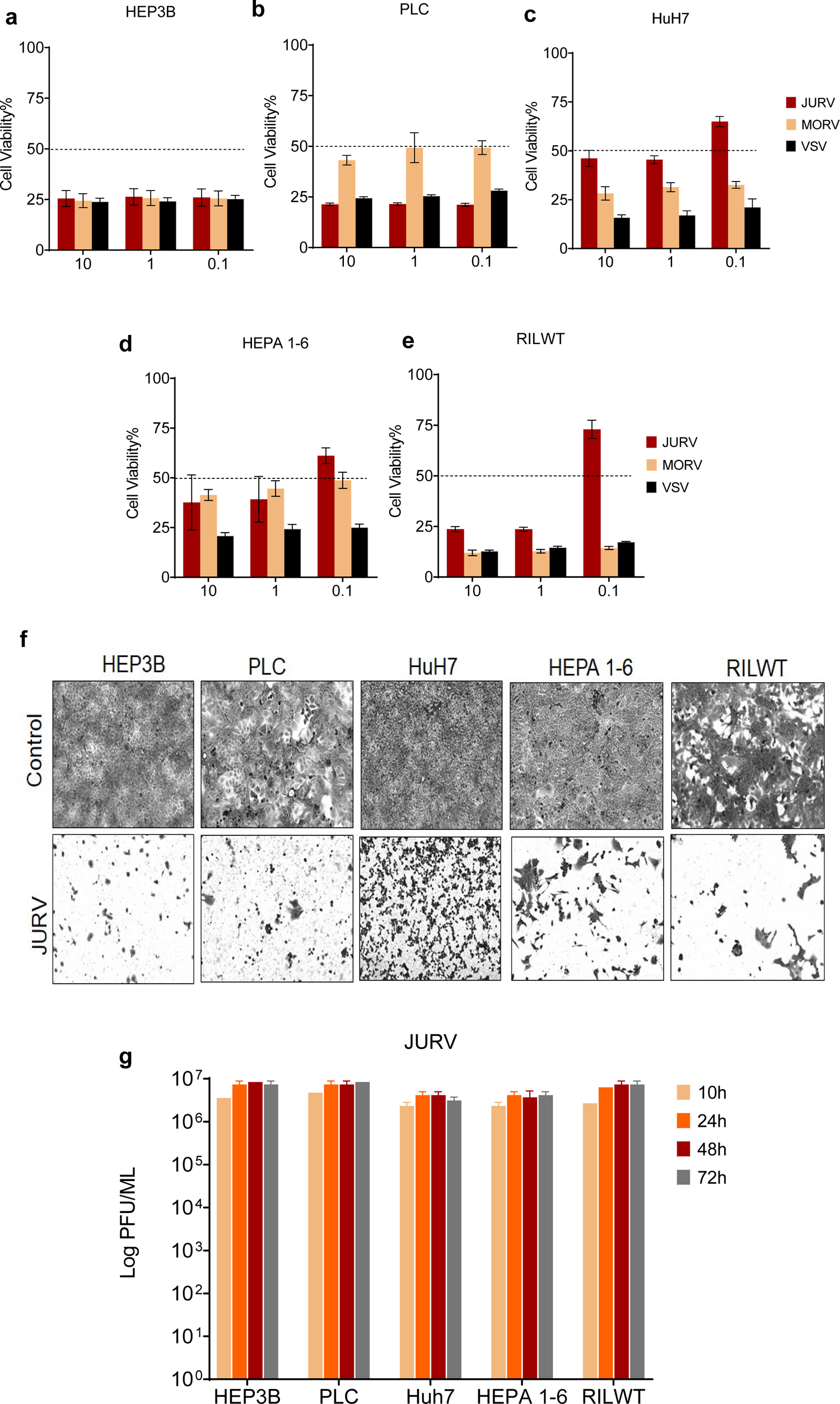
Oncolytic JURV is effective in inducing oncolysis in HCC cell lines. (**a**) The antigenome organization of jurona virus (JURV) consists of (3’ to 5’) of a leader (Le) sequence followed by approximately 10,993 bases comprising five structural genes (JURV-N, JURV-P, JURV-M, JURV-G, and JURV-L) separated by a highly conserved intergenic region (IGR) and a trailer sequence (Tr). Monolayers of human HCC, HEP3B (**b**), PLC (**c**), HuH7(**d**) and murine HCC, HEPA 1-6 (**e**) and RILWT(**f**) were seeded at a density of 1.5 x 10^4^/well in 96-well plates and infected with JURV, recombinant, morreton virus (MORV) and recombinant vesicular stomatitis virus (VSV) at the indicated multiplicity of infection (MOI) of 10, 1 and 0.1. The percentage of cell viability was determined 72 hours post-infection using a colorimetric assay (MTS, Promega USA). The discontinued lines on the graphs indicate the cut-off percentage for resistance (>50% cell viability above the line) and sensitivity (<50% of cell viability, below the line). Data was collected from multiple replicates over three independent experiments. Bars indicate mean ± SEM. (**g**) Crystal violet staining. Cancer cells were plated at 5.0 x 10^5^/well in a 6-well plate and rested overnight. The following day they were infected with JURV at an MOI of 0.1. Cells were fixed and stained with crystal violet 72 hours post-infection, and images were captured at 10x magnification on an Olympus IX83 Inverted Microscope System. (**h**) HCC cells were plated in 6-well plates at 2.0 x 10^5^/well and infected with JURV at MOI of 0.1. Supernatants from infected cells were collected at different time points, and viral titer was determined using a TCID_50_ (50% tissue culture infective dose) method on Vero cells (1.5 x10^4^). Data are plotted from two independent assessments of TCID_50_ for each point with mean ± SEM. (g) HCC cells (2.0 x10^4^) were pretreated with various concentrations of universal type I interferon-alpha (IFN-α), infected with viruses at an MOI of 0.01. Cell viability was measured using an MTS assay after 48 hours post-infection.

### Pretreatment with species-specific IFN-β does not protect HCC cells from virus-mediated tumor cell death

Antiviral response mediated by type I interferon (IFNs) constitutes an essential element of the innate immune system response to viruses.^7, 8^ Studies have shown defects in IFN signaling pathways that coincide with carcinogenesis create needful conditions for tumor-selective replication of oncolytic viruses (OVs).^9, 10^ In contrast, normal cells are protected because they possess functional antiviral responses that inhibit viral replication.^9^ It is now well established that the tumor microenvironment is a microcosm of complex interactions between tumorigenic and non-cancer cells.^11^ As a result, upon sensing the virus, non-cancer cells could potentially secrete antiviral cytokines (i.e., IFN-α, IFN-β) that can prematurely impair OVs replication, spread, and oncolytic activity if the cancer cells are responsive to the effect of exogenous IFNs.^12^ Therefore, to determine the impact of exogenous type I IFN on the outcome of JURV infection, we compared the susceptibility to JURV infection of monolayers of HCC cells (HEP3B, HEPA 1-6) mock-treated or pretreated with serial concentrations of species-specific IFN-β. Our data indicate that treatment with IFN-β did not shield (30-75% of cell death) human HCC (HEP3B) and mouse HCC (HEPA 1-6) cells from JURV-induced cell death (Supplemental Fig. 2 a, b). However, we noted a difference in response to IFN between the HEPA 1-6 and HEP3B cell lines. Indeed, at an MOI of 1, a concentration of 500 U/mL IFN-β killed approximately 20% of HEPA 1-6 (Supplemental Fig. 2 b), whereas the same treatment yielded ∼ 75 % of cell killing in HEP3B cells (Supplemental Fig. 2 a).

**Figure 2.**
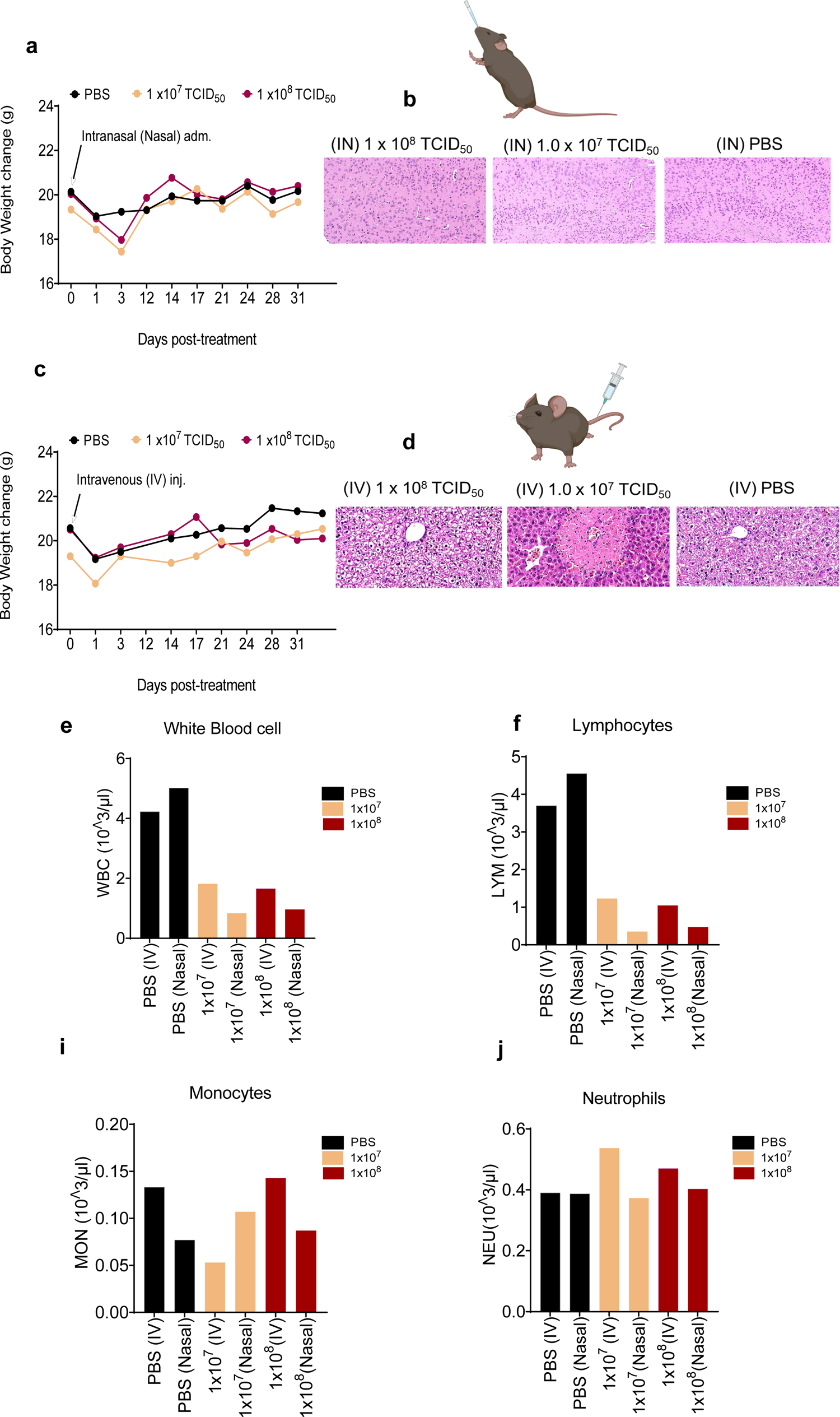
Effects of low and high doses of oncolytic JURV on body weight and hemogram in mice. Non-tumor bearing female C57BL6/J (n=6/group; Strain #:000664) of 6-8 weeks of age were administered intranasally (IN) or intravenously (IV) single doses of 1 x10^7^ TCID_50_ or 1 x10^8^ TCID_50_ of JURV. Body weight was recorded twice a week in both the IN and IV cohorts to assess drug-related toxicity. Three mice per group in each cohort (IN &IV) were sacrificed three days post-infection, and blood, brain, and liver were harvested to assess the short-term toxicity. Changes in body weight and hematoxylin and eosin (H&E) staining (brain and liver) are shown for mice administered IN (**a, b**) and IV (**c, d**) in mice treated with low (1 x10^7^ TCID_50_) and high low (1 x10^8^ TCID_50_) doses of JURV. Complete blood count, including white blood cells (**e**), lymphocytes (**f**), monocytes (**i**), and neutrophils (**j**), assessment of JURV-induced toxicity.

### High-dose intranasal or systemic virus challenge is not associated with clinically significant adverse events in non-tumor-bearing mice

To determine whether acute neurotoxicity or hepatoxicity can result from exposure to oncolytic JURV as shown for VSV^13–15^, non-tumor bearing mice were injected with single relatively low or high doses of attenuated oncolytic JURV. Reports have shown that intranasal (IN) administration of approximately 1 x 10^6^ TCID_50_ of wild-type VSV in mice results in significant weight loss and lethality at around three days post-infection.^16–18^ Here, we administered wild-type attenuated JURV to mice at a 10 to 100-fold higher dose (1 x 10^7^ − 1 x 10^8^ TCID_50_) intranasally, but also intravenously (IV) to mimic the natural route of systemic VSV infection. At three days post-infection, n=3/group were sacrificed, and blood, brain, liver, and spleen were collected to assess short-term toxicity. There was a slight drop in body weight (10-15%) on day 3 in both the intranasal (Fig. 2 a) and intravenous (Fig. 2 c) experiments. Histopathological image analysis of brain (Fig. 2 c) and liver sections of mice (Fig. 2 d, Supplemental Fig. 2 a) treated with the lowest or highest dose of JURV indicate that administration of JURV was not associated with brain damage or liver toxicity. Systemic VSV infection is characterized by temporarily leukopenia.^19^ Consistent with the earlier findings, we noted in the short-term toxicity cohort a decrease in the number of white blood cells and lymphocytes than monocytes and neutrophils in mice treated with JURV (Fig. 2 e-j). There was no significant difference in body weight or clinical signs (i.e., paralysis, death, matted fur) in mice challenged with JURV compared to PBS in the long-term observation group.

### JURV intranasal or intravenous infection in mice results in differentially expression of cellular proteins, but not significant neurotoxicity or hepatoxicity

Quantitative proteomics was used to compare protein changes in the brain and liver following injections with JURV. Non-tumor-bearing healthy mice were inoculated intranasally (IN) or intravenously (IV) with JURV. Three days post-infection (n=3/group), mice were sacrificed, and brains and liver were harvested to assess changes in protein expression changes by mass spectrometry. At a dose of 1 x 10^8^ TCID_50_ of JURV, we found that 60 differentially expressed proteins (DEPs) were upregulated (*P*-value < 0.055 and 2-fold change > 2) in the brain following IN infection (Fig. 3 c, d), and 87 DEPs were upregulated in the liver following IV injection (Fig. 3 c, d). We found that several of the 10 top DEPs upregulated in the brain, such as Stat1^20^, Ifit3^17^, and Lgals9^21^ play essential roles in the control of RNA virus infection by binding and directly regulating the functions of viral or cellular proteins with antiviral activity (Fig. 3 e). Similarly, the top 10 DEPs upregulated in the liver, such as Isg15^22^, Cmpk2^23^, Uba7^24^, exhibit key cellular antiviral functions (Fig. 3 f). Furthermore, Ingenuity Pathway Analysis (IPA) analysis of the DEPs in the brain (Fig. 3 g) and in the liver (Fig. 3 h) infected with JURV predicted activation of the interferon signaling, mitochondrial dysfunction, acute phase response signaling pathways which are all related to sensing and induction of innate and adaptive immune response to viral infection. Interestingly, we found that changes in the DEPs following IN or IV administration of 1 x 10^7^ TCID_50_ of JURV were like that of a 10-fold higher dose (1 x 10^8^ TCID_50_) of the virus (Supplemental Fig. 3 a-d). Moreover, analysis of the top 10 DEPs upregulated in the brain (Supplemental Fig. 3 e) and liver (Supplemental Fig. 3 g) and predicted activated pathways (IPA) are all immune responses to viral infection-related proteins and pathways (Supplemental Fig. 3 g-h). These results indicate that JURV infection was sensed by the host cellular apparatus leading to the expression of proteins/cytokines with potent antiviral properties to eliminate the viral infection. In addition, it shows that injections of high doses of JURV did not elicit severe toxicity or result in physical impairment in the mice susceptible to VSV induced neurotoxicity.

**Figure 3.**
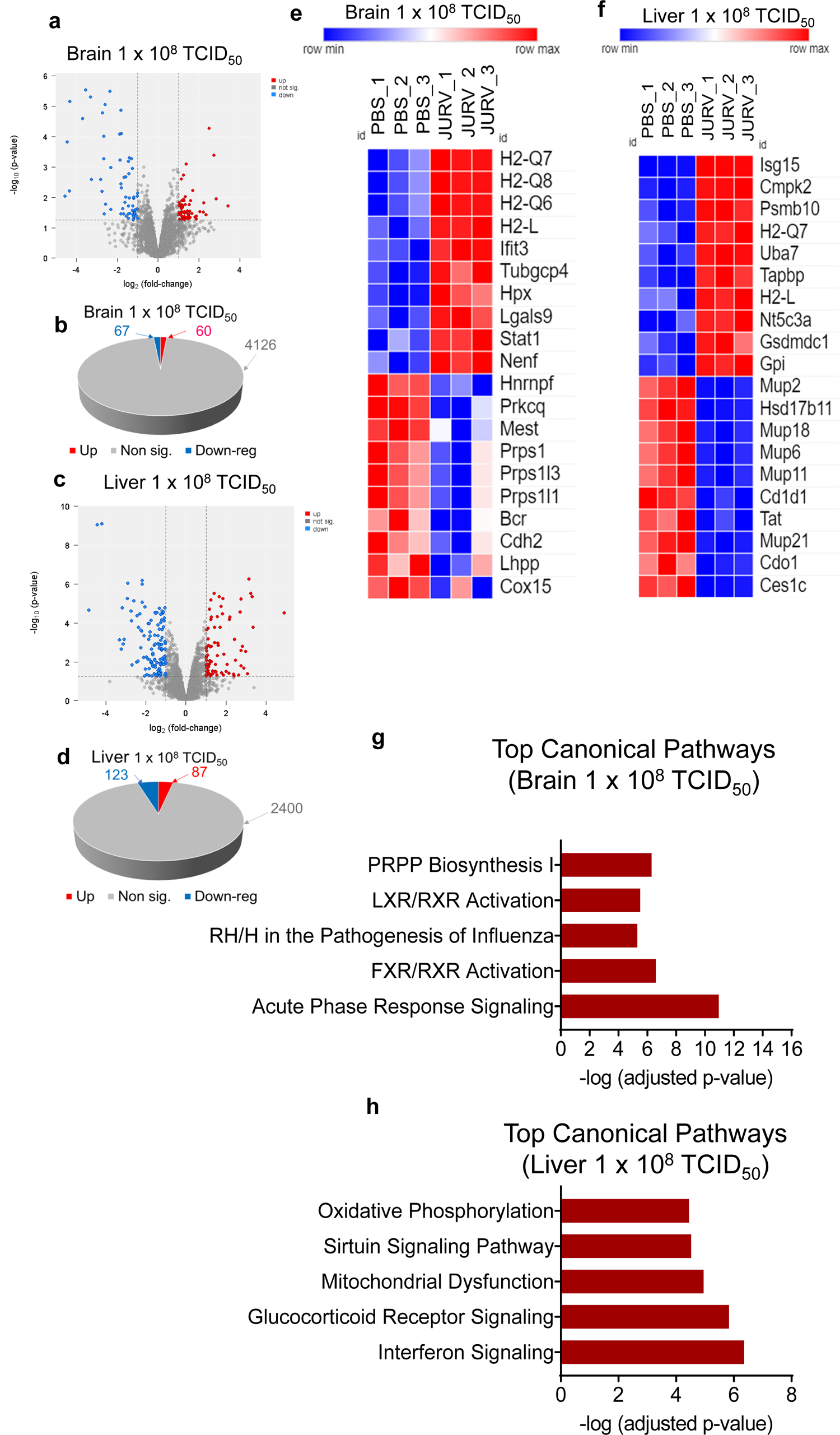
Toxicoproteomic analysis of brain and liver tissues of mice injected with high doses of oncolytic JURV. Intranasal and intravenous administration of oncolytic JURV in mice causes changes in the expression of proteins associated with sensing RNA viruses and antiviral signaling pathways. Volcano plot of protein expression differences for the brain (**a**) and liver (**c**) of mice treated with PBS vs. 1 x10^8^ TCID_50_ of JURV. 3-D Pie slices of the numbers of differentially expressed proteins (DEPs) for the brain (**b**) and liver (**d**) of mice injected with PBS vs. 1 x10^8^ TCID_50_ of JURV. Heatmaps of the top 20 DEPs up or down regulated in the brain (**e**) and liver (**f**) of mice injected with PBS vs. 1 x10^8^ TCID_50_ of JURV. DEPs were determined using the limma-voom method. A fold-change |*log*FC| ≥ 1 and a false discovery rate (FDR) of 0.055 were used as a cutoff. The *log*FC was computed using the difference between the mean of *log2*(JURV) and the mean of *log2*(PBS), that is, mean of *log-2*(JURV) - mean of *log2*(PBS). Graph showing top-scoring canonical pathways significantly enriched by treatment with 1 x10^8^ TCID_50_ of JURV in the brain (**g**) and liver (**h**).

### Intratumoral injections of JURV control local and distant tumor growth with an anti-tumor efficacy equal to that of anti-PD-1 therapy

Studies employing murine models of HCC have demonstrated improved anti-tumor efficacy when blocking the programmed cell death protein 1 (PD-1) ^25^ axis pathway compared to other drugs such as multikinase inhibitors (i.e., sorafenib); we then tested whether JURV co-administered with anti-PD-1 antibodies could yield a significantly greater anti-tumor efficacy than JURV alone. To do this, we injected intratumorally (IT) three doses of JURV alone, or anti-PD-1 alone or JURV combined with -PD-1 antibodies into the right flanks of subcutaneous mouse models of HCC (HEPA 1-6). Our data clearly show IT administration of JURV provides therapeutic efficacy as manifested by a prominent (***) tumor growth delay in the JURV-treated mice compared to the control group (PBS) (Fig. 4 a). The therapeutic efficacy of intraperitoneal (IT) injection (Fig. 4 b) of anti-PD-1 therapy was comparable (***) to that of JURV. However, the effectiveness of JURV combined with anti-PD-1 antibodies was less striking (**) than that of JURV or anti-PD-1 alone. Analysis of the individual tumor volumes of mice indicates that nearly all (>70%) mice in the JURV-treated and anti-PD-1-treated groups eliminated their tumors compared to JURV + anti-PD-1 antibodies (>50%) groups (Supplemental Fig. 4 a). We next sought to investigate whether local administration of JURV could elicit a systemic anti-tumor effect capable of not only impact local tumor development but also distant tumors. We subcutaneously implanted HEPA 1-6 tumors on the right and left flanks of immunocompetent mice; however, we performed local IT injections of JURV on the right flanks only, leaving the left flanks unaffected. When comparing the injected and non-injected flanks, we found that IT injections of JURV triggered an anti-tumor activity that reduced the growth of HEPA 1-6 tumors on both right and left flanks (Fig. 4 c, Supplemental Fig. 4 b). There was a drop in body weight a day post-injection of JURV, mice quickly recovered, and no adverse events were observed until the end of the study (day 28) (Supplemental Fig. 4 c). Although one mouse was found dead in the control (PBS) group, there was no difference in survival between the different groups (Supplemental Fig. 4 d). Moreover, an assessment of the level of treatment-associated liver and kidney toxicity biomarkers in the serum indicates that mice did not experience severe virus-induced toxicity (Supplemental Fig. 5 a-o). This finding aligns with the induction of systemic anti-tumor activity commonly observed with oncolytic viruses in clinical trials and animal models.^3,26^ Taken together, this data highlights the potential of JURV to induce robust anti-tumor activity against local and distant non-injected tumors, which is critical for targeting oligometastatic or metastatic diseases.

**Figure 4.**
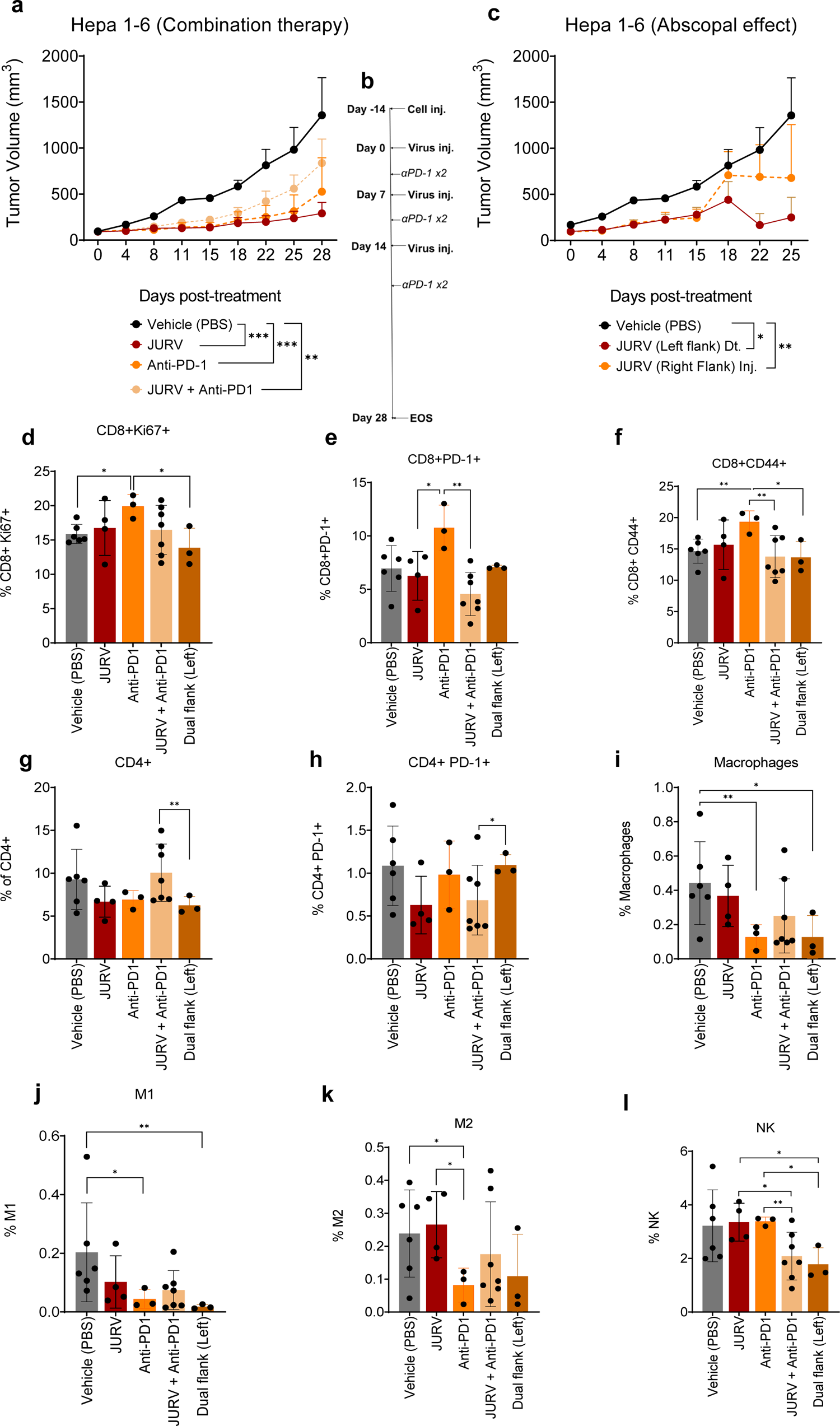
Evaluation of the anti-tumor efficacy of oncolytic JURV as monotherapy or combined with anti-PD-1 antibodies in murine HCC model. HEPA 1-6 cells were implanted into the right flanks of female C57BL6/J (Strain #:000664) (n=7/group; Jackson Laboratory). (**a**) Upon reaching 80-120 mm^3^, mice were administered 50 µL IT injections containing PBS (Vehicle), 1 x 10^7^ TCID_50_ units of JURV, anti-PD-1 therapy, or combination JURV + anti-PD-1. Anti-PD-1 antibodies (10mg/kg) were given intraperitoneally (IP) twice a week for three weeks. (**b**) In the treatment schedule, at day −14, tumor cells were subcutaneously implanted into the right flanks of mice, then PBS, JURV, anti-PD-1 antibodies (αPD-1), JURV + anti-PD-1 was injected (inj.) into tumor-bearing mice at days 0, 7, and 14. Tumors were harvested at the end of the study for downstream analysis. (**c**) In the abscopal model (dual flanks), HEPA 1-6 cells (1 x 10^6^ cells/mouse) were first subcutaneously grafted into the right flanks and were categorized as “primary” tumors. Simultaneously, we performed distant HEPA 1-6 tumor grafts injection (1 x 10^6^ cells/mouse) into the left flanks of these mice. Mice in the dual flank group received 50 µL IT injections of 1 x 10^7^ TCID_50_ units of JURV on their right flanks only once a week for three weeks. Data plotted as mean +/- SD, **p<0.001 ***p<0.0001. Area under tumor growth curves was compared by one-way ANOVA with Holm-Sidak correction for type I error. The day when we injected the first JURV or PBS into the mice is defined as day 0. Combination JURV + anti-PD-1 therapy profoundly modulated tumor microenvironment as manifested by changes in the frequency of (**d**) CD8+ Ki67+, (**e**) CD8+PD-1+, (**f**) CD8+CD44+, (**g**) CD4+, (**h**) CD4+ PD-1+, (**i**) macrophages, (**j**) M1-like macrophages, (**k**) M2-like macrophages, and (**l**) NK cells. The sub-myeloid population representing macrophages are CD11b+F480+ cells, and the two markers used to segregate the M1 and M2 sub-populations are CD206 and I-A/I-E, known as major histocompatibility class II (MHCII) in mouse. The Bartlett test was used to test homogeneity of variance and normality. If the p value of the Bartlett test was no less than *P*=0.05, ANOVA and a two-sample t test were used to compare group means. If the *P* value of the Bartlett test was less than 0.05, the Kruskal-Wallis and Wilcoxon rank-sum tests were used to compare group means. The figures demonstrate the potential significant difference of the gated subsets in the CD45+ population, determined by the p value.

**Figure 5.**
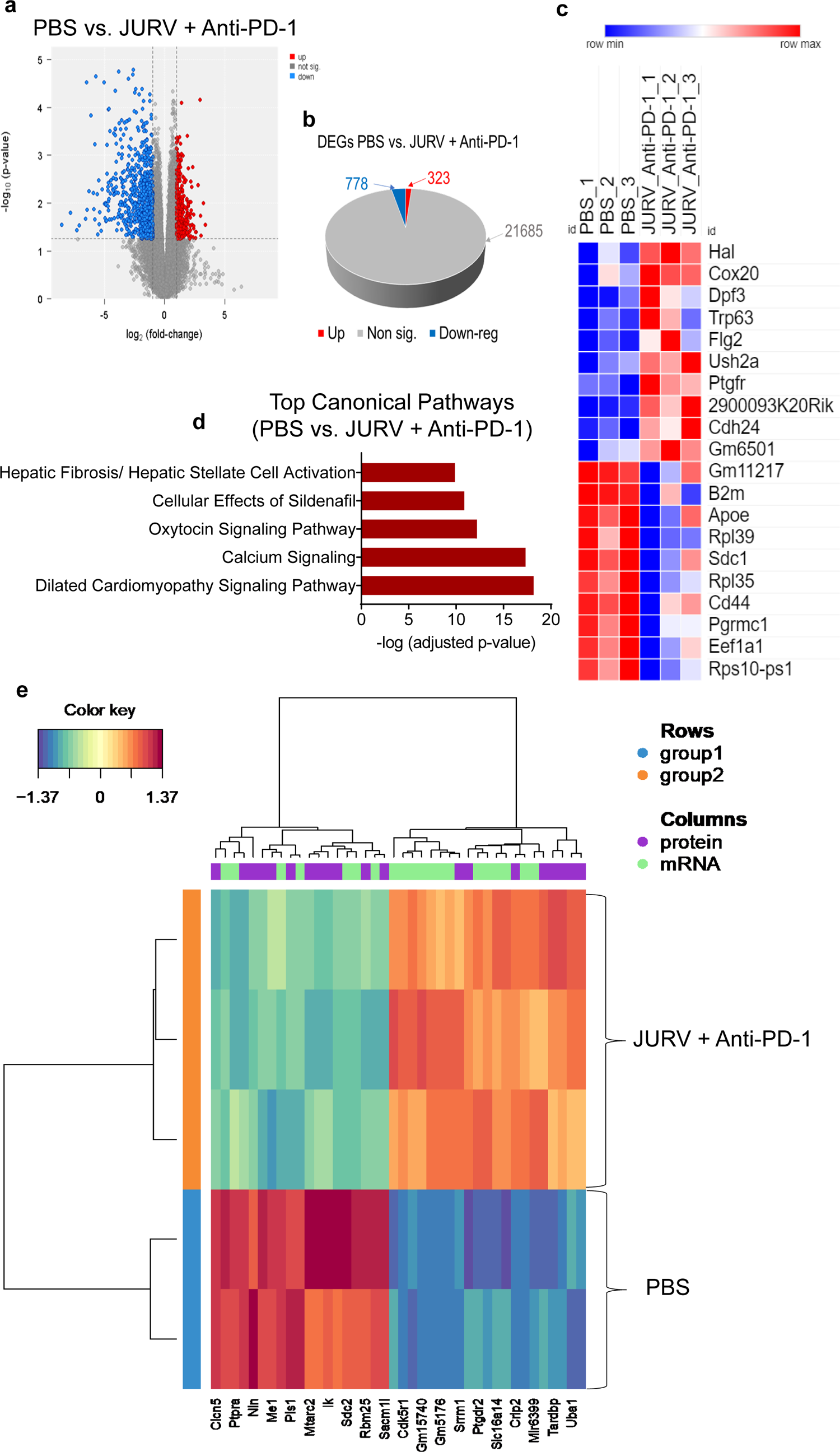
Proteogenomic response of murine HCC to combination oncolytic JURV and anti-PD-1 therapy. HEPA 1-6 tumors were harvested from mice treated with PBS or JURV + anti-PD-1 antibodies. (**a**) Volcano plot of mRNA expression differences for PBS vs. JURV (1 x10^8^ TCID_50_ x 3) + anti-PD-1 antibodies (10mg/kg x 6). (**b**) 3-D Pie slices of the numbers of differentially expressed genes (DEGs) between PBS vs. JURV + anti-PD-1 antibodies. (**c**) Heatmap of the top 20 DEGs up- or down-regulated PBS vs. JURV + anti-PD-1 antibodies. DEGs were determined using the limma-voom method. A fold-change |*log*FC| ≥ 1 and a false discovery rate (FDR) of 0.055 were used as a cutoff. The *log*FC was computed using the difference between the mean of *log2*(JURV + anti-PD-1) and the mean of *log2*(PBS), that is, mean of *log2*(JURV + anti-PD-1) - mean of *log2*(PBS). (**d**) Graph showing top-scoring canonical pathways significantly enriched by treatment with JURV + anti-PD-1 antibodies. A MixOmics supervised analysis was carried out between differentially expressed proteins (DEPs) and DEGs based on Log2 fold change values. Log2 fold change of DEG × Log2 fold change of DEP > 0 with a *P-value* of DEG and DEP <0.05 were considered associated DEGs/DEPs. (**e**) DEG/DEP expression heatmap of the 30 most up-regulated and down-regulated features DEG/DEP in JURV + anti-PD-1 antibodies vs. PBS.

### PD-1 blockade combined with IT injection of JURV profoundly modulates the immune component of the tumor microenvironment

Many studies have shown that vesiculoviruses selectively infect, replicate, and lyse tumor cells and modulate local and systemic anti-tumor immune responses.^27^ Given that administration of JURV, anti-PD-1 antibodies or a combination of JURV and anti-PD-1 antibodies evoked prominent tumor regressions in the subcutaneous syngeneic HCC model. Therefore, we sought to investigate and compare the changes in the immune responses associated with these different treatment regimens in mice bearing dual tumors (left flanks). As expected, our data show dramatic changes in the frequencies of tumor-infiltrating lymphocytes (TILs) after administering these therapies to mice. When comparing the treatment groups, we observed that IT injection of JURV significantly decreased the subset of (**) F4/80-TILs (Supplemental Fig. 6 b) and increased the distribution of (*) M2-like macrophages (Fig. 4 k), (*) natural killer (NK) cells (Fig. 4 l), and (*) natural killer T (NKT) cells (Supplemental Fig. 6 a) compared to control (PBS), anti-PD-1, and dual flank (left). Analogously, immune blockade of PD-1 correlated with the intratumoral accumulation of cytotoxic CD8+ T cells, mainly (*) CD8+ Ki67+ (Fig.4 d), (*) CD8+ PD-1+ (Fig.4 e), and (**) CD8+ CD44+ (Fig.4 f), compared to PBS, JURV, and dual flank (left). We also noted a prominent decrease in the TILs macrophages (**) (anti-PD-1 vs. PBS) (Fig.4 i), (*) M1 (anti-PD-1 vs. PBS) (Fig.4 j), and (*) M2-like (anti-PD-1 vs. JURV) (Fig.4 k), in the anti-PD-1 treated group. These results show that IT treatments with JURV elicit anti-tumor immunity via recruitment and tumor infiltration of cytotoxic effectors T (CTLs) cells.^28, 29^ We also observed an increase in M2-like macrophages, which along with myeloid cells, are known to disable anti-tumor immunity via expression and interaction of PD-L1 with PD-1 on the surface of cytotoxic T cells.^30, 31^ Thus, unsurprisingly, we found that combining JURV and anti-PD-1 antibodies profoundly modulate both adaptive and innate immunity as manifested by an increase in (**) CD4+ (Fig. 4 g), (*) F4/80-(Supplemental Fig. 6 b), (*) CD4+ PD-1+ (Fig. 4 h), (*) CD11b+ (Supplemental Fig. 6 d), (**) CD11b-(Supplemental Fig. 6 e), and decrease in (**) CD8+ PD-1+ (Fig. 4 e), (*) (**) NK (Fig. 4. l) compared to JURV, anti-PD-1, and dual flank (left). No significant change was observed in serum IFN-β in mice treated with JURV, anti-PD-1, and JURV combined with anti-PD-1 antibodies and dual flanks (Supplemental Fig. 7 a). Collectively, our data suggest that the combination of JURV and anti-PD-1 antibodies primes robust anti-tumor immunity by increasing cytotoxic TILs in the tumor and inhibiting immunosuppression, the hallmark of durable response to immunotherapy.

**Figure 6.**
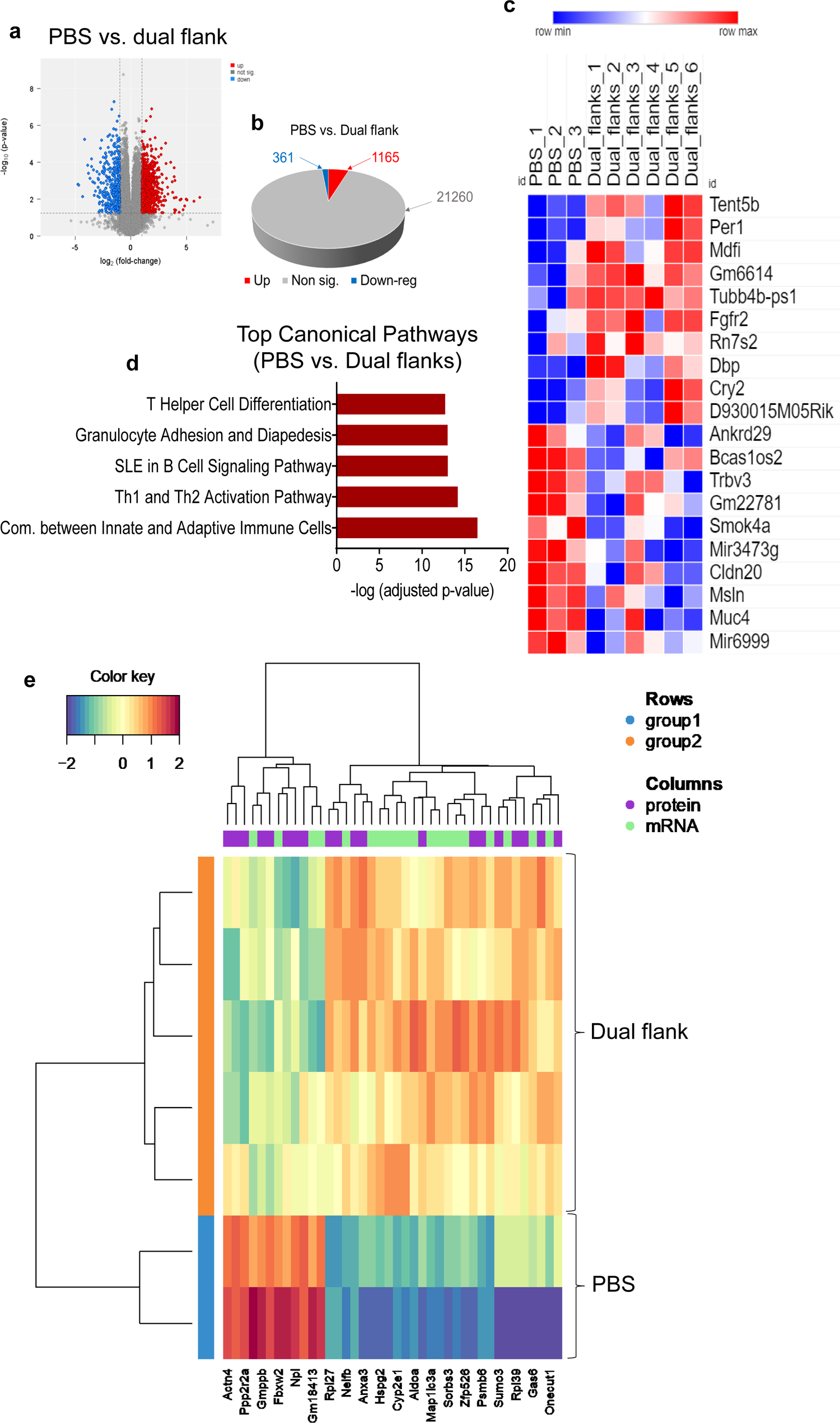
Deciphering the mechanisms of oncolytic JURV mediated anti-tumor activity in local and distant murine HCC. We performed transcriptomic analysis on HEPA 1-6 tumors harvested from the non-injected HEPA 1-6 tumors in the abscopal effect cohort. (**a**) Volcano plot of mRNA expression differences for PBS vs. dual flanks (non-injected tumors). (**b**) 3-D Pie slices of the numbers of differentially expressed genes (DEGs) between PBS vs. dual flanks (non-injected tumors). (**c**) Heatmap of the top 20 DEGs up or down regulated PBS vs. dual flanks (non-injected tumors). DEGs were determined using the limma-voom method as described in Fig. 5. (**d**) Graph showing top-scoring canonical pathways significantly enriched by treatment with PBS vs. dual flanks (non-injected tumors). A MixOmics supervised analysis was carried out between DEPs and DEGs based on Log2 fold change values. Log2 fold change of DEG × Log2 fold change of DEP > 0 with a *P-value* of DEG and DEP <0.05 were considered associated DEGs/DEPs. (**e**) DEG/DEP expression heatmap of the 30 most up-regulated and down-regulated features DEG/DEP in PBS vs. dual flanks (non-injected tumors).

**Figure 7.**
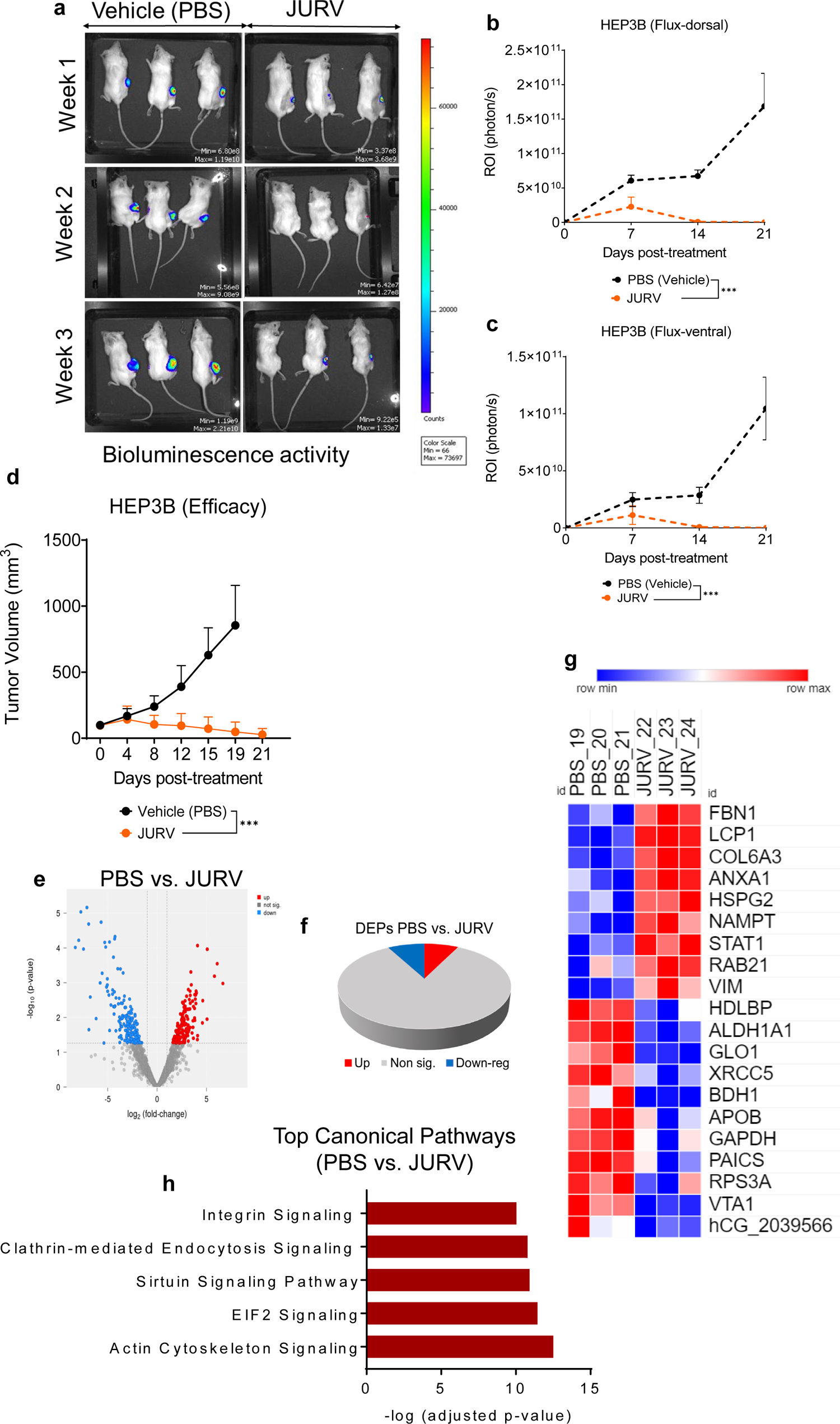
Oncolytic JURV induces a robust virus-mediated oncolysis dependent tumor growth delay in JURV in Hep3B Xenografts. Female NOD.Cg-*Prkdc^scid^*/J (Strain #:001303) mice (n=6/group) were inoculated subcutaneously with human luciferase expressing HEP3B cells. (**a**) Photon emission from mice with subcutaneous HEP3B tumors. When average tumor volume was over 80-120 mm^3^, mice were divided into two groups and received IT injections with either PBS, JURV at a dose of 1 x 10^7^ TCID_50_ on days 0, 7 and 14. Photon counts were obtained on weeks 1, 2, and 3. Photon emission at the right dorsal flanks (**b**) and ventral (**c**) were significantly (*P*<005) reduced among JURV-treated mice than in PBS. (**d**) A digital caliper was used to determine tumor volume twice weekly until a humane endpoint, death, or end of study (Day 21). Data plotted as mean +/- SD. Paired t test with and Wilcoxen signed-rank test were performed (95% CI), ***P<0.005. HEP3B tumor treated with PBS or JURV was harvested, and we analyzed changes in protein expression. (**e**) Volcano plot of protein expression differences in HEP3B tumors treated with PBS vs. 1 x10^7^ TCID_50_ of JURV. (**f**) 3-D Pie slices of the numbers of differentially expressed proteins (DEPs) in HEP3B tumors injected with PBS vs. 1 x10^7^ TCID_50_ of JURV. (**g**) Heatmap of the top 20 DEPs up or down regulated in HEP3B tumors injected with PBS vs. 1 x10^7^ TCID_50_ of JURV. DEPs were determined using the limma-voom method as described in Fig 3. A fold-change |*log*FC| ≥ 1 and a false discovery rate (FDR) of 0.055 were used as a cutoff. The *log*FC was computed using the difference between the mean of *log2*(JURV) and the mean of *log2*(PBS), that is, mean of *log2*(JURV) - mean of *log2*(PBS). **(h)** Graph showing top-scoring canonical pathways significantly enriched by treatment with 1 x10^7^ TCID_50_ of JURV in the HEP3B tumors.

### Analysis of associated DEGs/DEPs following administration of JURV in murine HCC identified pathways involved in inflammation and innate and adaptive immune responses

To determine the impact of IT administration of JURV on gene expression profiles in the tumor, we examined the transcriptome of murine HCC tumors injected with three doses of JURV. Differentially expressed genes (DEGs) were analyzed using the limma-voom method.^32^ Our data (Supplemental Fig. 8 a, b) showed that 203 genes were up-regulated and 463 genes were downregulated (*p*<0.055). Several of the 10 up-regulated DEGs Myo3a^33^, Cd209c^34^, Trim67^35^, St8sia2^36^, and Wnt5b^37^ were associated with the immune response pathways (Supplemental Fig. 8 c). Moreover, IPA was used to identify enriched cellular signaling pathways associated with gene expression changes following IT administration of JURV. Many of these pathways, such as the B Cell Receptor Signaling, IL-15 Signaling, and Phagosome formation, are related to activation of the host innate and adaptive immune responses (Supplemental Fig. 8 d). Furthermore, to comprehensively examine the mechanism of JURV-induced anti-tumor activity, we analyzed the DEPs and DEGs from the transcriptomic and proteomic data (Supplementary Fig. 8 e). In the associated DEGs/DEPs, we identified the top 30 enriched features that were significantly upregulated or downregulated in the JURV group compared to the control group (PBS). Among the upregulated features, the S1pr3^38^, Tnpo1^39^, Psmb10^40^, Ddt^41^, Ncor2^42^, Slc04c1^43^ have been identified in inflammation, host immune response against microorganisms (virus, bacteria) and tumorigenesis.

### PD-1 blockade in murine HCC modulates inflammation while enhancing anti-tumor immunity

Immune checkpoint blockade employing PD-1 inhibitors or B7-cytotoxic T-lymphocyte antigen 4 (CTLA-4) inhibitors improve CTLs (i.e., CD8+) anti-tumor immune responses, tumor growth suppression, and survival in clinical trials and animal model of HCC.^44, 45^ Our data show that IP administrations of anti-PD-1 antibodies are accompanied by a robust tumor regression and effectively establish anti-tumor immunity in HEPA 1-6 tumor-bearing mice. Therefore, here, we aimed to decipher the transcriptional and proteomic changes that happened upon IP treatments with anti-PD-1 antibodies. When scrutinizing mRNA expression profiles in murine HCC tumors, we found that 860 genes were up regulated and 241genes were downregulated in the HEPA 1-6 tumors (Supplementary Fig. 9 a, b). All the top 10 DEGs (Supplementary Fig. 9c), including Prps1l1^46^, Cstdc4^47^, and Ppbp^48^ that were upregulated in the anti-PD-1 therapy group compared to control (PBS), are involved in inflammation, adaptive immune response, and cell survival and proliferation. Similarly, most enriched pathways (Supplementary Fig. 9 d) predicted by IPA analysis (i.e., Phagosome, communication between Innate and Adaptive Immune Cells) are immune-related pathways. Integrated analysis of the associated DEPs/DEGs features (Supplementary Fig. 9 e) revealed upregulation of Tmod3^49^, Nfkb2^50^, Dapk3^51^, Nipsnap3b^52^, Sart3^53^, and Adgrl4^54^ are key effectors of pathways regulating immunosuppression, angiogenesis, inflammation and anti-tumor immunity.

### Proteogenomics changes following IT treatments with the combination of JURV, and anti-PD-1 antibodies correlate with activation of innate and adaptive immunity

We analyzed the transcriptional profile of murine HCC to identify genes and pathways dysregulated in tumor following treatment with JURV, anti-PD-1 antibodies, and combination JURV and anti-PD-1 antibodies as compared to control (PBS). While comparing DEGs between the control group (PBS) and combination JURV and anti-PD-1 antibodies, we found that 323 genes were up-regulated, whereas 778 genes were downregulated (Fig. 5 a, b). The top 10 up regulated DEGs, including the Cox20^55^, Dpf3^56^, Trp63^57^, Flg2^58^, Ush2a^59^, and Cdh24^60^, are effector genes associated with tumor suppression and immune cells infiltration mechanism (Fig. 5 c). IPA analysis identified enriched pathways (i.e., Hepatic Fibrosis/Hepatic Stellate Cell Activation, Oxytocin Signaling, Calcium Signaling pathways) which are all primarily associated with activation, and regulation of immune cells (Fig. 5d). Integrated analysis of the associated DEPs/DEGs (Fig. 5 e), highlighted the up-regulation of genes linked to susceptibility to immunotherapy, autophagy, tumor repression and immune responses such as Cdk5r1^61^, Ptgdr2^62^, Crip2^63^, Tardnp^64^ (Fig. 5 e).

Furthermore, examination of unique and common DEGs between PBS vs. JURV, PBS vs. Anti-PD-1, and PBS vs. JURV + Anti-PD-1 (Supplemental Fig. 10 a) uncovered a relatively small overlap between co-differentially expressed genes (both up-regulated and down-regulated) among these three groups, suggesting the existence of substantial differences in the mechanism of anti-tumor responses to these therapies, potentially driven by differences in the type of TILs these therapies activate, recruit and traffic to the tumor site or between the two routes of delivery (IP and IT). Moreover, we performed gene ontology (GO) term enrichment analysis on 35 up-regulated or down-regulated DEGs (Supplemental Fig. 10 b) common to all three data sets. Some mitogen-activated protein kinase (MAPK) ^65^ pathways known to be up-regulated during virus infection, macrophage migration inhibitory factor (MIF) ^66^, involved in inflammatory and immune response, necroptosis signaling, and vascular endothelial growth factor (VEGF) family were found to be enriched (Supplemental Fig. 11 a). These data indicate that even though JURV and anti-PD-1 therapies display different mechanisms of anti-tumor activity, they activate important and complementary pathways involved in innate and adaptive anti-tumor immunity.

### Administration of JURV activates molecular mechanism involved in anti-tumor immunity that targets both local and distant murine tumors

Previous reports have demonstrated that oncolytic viruses can induce virus-mediated activation of local and systemic anti-tumor immunity through the recruitment of class I MHC-restricted virus-specific and tumor-specific CTLs.^67, 68^ To better comprehend the mechanisms behind the observed abscopal effect of JURV on distant tumors, we analyzed the transcriptome and proteome of non-injected (left flank) HEPA 1-6 tumors. Comparison of DEGs between the control group (PBS) and dual flank (left flank) showed that 1165 genes were up-regulated, and 361 genes were down-regulated (Fig. 6 a, b). Several top 10 DEGs, such as Tent5b^69^, Per1^70^, Tubb4b-ps1^71^ and fgfr2^72^ were immune-related genes associated with activation of humoral immune responses, regulation of excessive immune responses, tumor invasiveness, immune process, and circulatory system (Fig. 6 c). Consequently, most enriched pathways (i.e., T Helper Cell Differentiation, Th1, and Th2 Activation, and Communication between Innate and Adaptative Immune cells) identified by IPA analysis suggest that modulation of the T helper cell pathways^73^ played a crucial role in the abscopal effect induced by IT injection of JURV (Fig. 6 d).

A comparison of DEPs/DEGs illustrates that most up-regulated features, including Anxa3^74^, Hspg2^75^, Cyp2e1^76^ and Map1lc3a^77^ are therapeutic targets in immunotherapy or function as tumor suppressor genes (Fig. 6 e). In addition, to assess the relationship between the local and systemic anti-tumor immunity in the murine tumors, we performed Go term enrichment analysis of DEGs between the JURV-injected and non-injected HEPA 1-6 tumors. The top enriched canonical pathways (i.e., Inhibition of ARE-Mediated mRNA Degradation, FAT10 Signaling, EILF2 Signaling) identified by analysis of DEGs between JURV vs. Dual flank are linked to response to immunotherapy, activation of MAPK pathways, and apoptosis (Supplemental Fig. 12a). These results indicate that JURV can effectively prime anti-tumor immunity against local and distant tumors in an animal model of HCC.

### IT administration of JURV mediates robust anti-tumor efficacy HEP3B-xenograft Models of HCC

We have previously shown that responsiveness to I IFN production or viral kinetic *in vitro* by infected cancer cell lines does not always correlate with in vivo efficacy to oncolytic.^4^ To determine whether JURV can induce an oncolysis-dependent tumor growth delay, we injected IT three doses of JURV into the immunocompromised HEP3B-xenografts. We employed luciferase tagged HEP3B cells to monitor tumor growth during the first three weeks of treatment. Compared to the control group (PBS), bioluminescence imaging showed a remarkable (***) tumor inhibition (Fig. 7 a-c) in the JURV-treated mice, which was visible from the first-week post-injection to the end of the study. In addition, to luciferase activity, a comparison of tumor volumes between PBS vs. JURV indicates that JURV triggered a significant (***) tumor growth delay (Fig. 7 d, Supplemental Fig. 13 a). We next perform proteomic to determine the changes occurring after IT deliver of JURV in HEP3B tumors. Analysis of DEPs showed a storm of 860 up-regulated proteins and 241 down-regulated proteins (Fig. 7 e, f). Out of the 10 top up-regulated DEPs, we identified LCP1^78^, COL6A3^79^, HSPG2^80^, NAMPT^81^, STAT1^82^ and VIM^83^ as proteins associated with activation of the mTORC2/AKT pathways, prognostic and treatment of cancer, regulation of tumor growth, and cancer cell stiffness. There was a drop (∼15%) in the body weight in the JURV group starting on day 19 (Supplemental Fig. 13 b). However, one mouse was found moribund on day 18, which we attributed to an isolated adverse due to the severe immunodeficient nature^84^ of the NOD-SCID mouse model as no other mice experienced toxicity (Supplemental Fig. 13 c). Nonetheless, our findings show that JURV induces a potent oncolysis in vitro and in vivo in human HEP3B-xenografts, which further increases enthusiasm around using the new immunovirotherapy in human HCC.

## DISCUSSION

Rhabdoviruses possess several advantageous properties over other oncolytic viral vector platforms, including amenability to genetic manipulation, and not using humans as a natural host resulting in low seroprevalence in the population.^5^ In addition to an episomal^85^ and fast kinetic cycle in tumor cells, most vesiculoviruses can encode large transgenes^4^ while maintaining the ability to potently infect, replicate and induce apoptosis^86, 87^ in a vast array of cancer cells.^88–94^ We and others have shown that others non-vesicular stomatitis virus (VSV) members^4, 95^ of the *Rhabdoviridae* family have tremendous potential as oncolytic agents because of their natural tumor selectivity and hypersensitivity to type 1 interferon response which is defective in 80% of tumors.^6, 7^

We described the anti-cancer potential of a laboratory attenuated Jurona virus (JURV), which displayed all the characteristics required for a potent immunovirotherapy, such as strong cytolytic effect and rapid replication cycle in tumor cells, sensitivity to type I IFN, and, more importantly, lack of long-term neurotoxicity and hepatotoxicity in mice. Furthermore, we showed that intratumoral administration of JURV induces oncolysis-mediated tumor regression in human HCC xenografts and elicits anti-tumor immunity via recruitment and activation of cytotoxic T (CTLs) cells leading to tumor growth delay in both local and distant murine tumors in a syngeneic HCC model. Moreover, when administered concomitantly, JURV and anti-PD-1 therapy considerably modulate the TME via enhanced infiltration of cytotoxic T-cells. However, we found that JURV and anti-PD-1 antibodies activate different effectors of the immune system, they have complementary anti-tumor activities. This suggests an attractive strategy in which, first, IT injections of JURV will be used to turn “cold” tumors into “hot” tumors, followed by treatment with immune checkpoint inhibitors to amplify the immune response and overcome immunosuppression for possible additive or synergistic long-term responses.

Analysis of mRNA and protein expression profiles in HCC tumors following administration of JURV, anti-PD-1 antibodies, and combination JURV and anti-PD-1 therapy unveiled the up-regulation of immune-related genes and identified enriched pathways involved in inflammation, regulation of immunosuppression, angiogenesis, and anti-tumor immunity. Our proteogenomic data further predicted that the T helper cell pathways might be a significant player in the JURV-induced abscopal effect.

Together, our results demonstrate that JURV potently infect HCC cells and induce oncolysis *in vitro* and in animal model. It also indicates that JURV-infected tumor cells prime an anti-tumor immunity (targeting primary and distant tumors) which is enhanced by addition of anti-PD-1 antibodies. These initial findings will hopefully support further development of this promising new oncolytic vector as a novel anti-neoplastic therapy in hepatocellular carcinoma, a disease with the most pressing clinical need.

## METHODS

### Experimental design

These experiments were performed to provide new and critical mechanistic insights into the safety and the efficacy oncolytic JURV in HCC tumor models, which will enable the rational design of studies using JURV as monotherapy or conjugated with other cancer therapies in early- or late-stage HCC for possibly additive or perhaps synergistic long-term responses in clinical settings. All animals were randomly allocated to the different study groups in an unblinded fashion. Average tumor volume (mm^3^) for each group + SEM at randomization was set between 80-120 mm^3^. The tumor volume (or its log transformation) was assessed in its relationship to time through a mixed linear regression model, using time, treatment effect, and their interaction as the independent variables. We used a random effect to account for within-subject correlation due to repeated measurements. Slopes were interpreted as growth rates (or logarithm) of the tumor over time and compared between groups. Analysis of Kaplan-Meier curves was used to identify the proportion of tumor-bearing mice living for a specific time after treatment. The *n* values and statistical methods are indicated in the statistical analysis section.

### Cell lines

This study used a panel of three human hepatocellular carcinomas (HCC) cell lines (HEP3B, PLC, HuH7) and two murine HCC cell lines (HEPA 1-6, R1LWT). All cell lines were cultured at 37° C with 5% CO2 in media supplemented with antibiotic agents (100 µg ml-1 penicillin and 100 µg ml-1 streptomycin). HEP3B, PLC, and HuH7 were maintained in Dulbecco’s Modified Eagle’s Medium (DMEM) with 10% fetal bovine serum (FBS). We also maintained HEPA 1-6, RILWT, BHK-21 (Baby Hamster kidney fibroblast), and Vero (African green monkey kidney) cells in DMEM with 10% fetal bovine serum (FBS). BHK-21 and Vero cells were obtained from the American Type Culture Collection (Manassas, VA). We purchased HEP3B, PLC, HuH7 and HEPA 1-6 from the American Type Culture Collection (ATCC, Manassas, VA). RILWT cell line was a gift from Dr. Dan G. Duda at MGH, Boston, MA.

### Oncolytic viruses

We obtained jurona virus (JURV) from the University of Texas Medical Branch (UTMB) World Reference Center for Emerging Viruses and Arboviruses (WRCEVA). A laboratory-adapted viral clone of JURV was generated using sequential plaque purifications on Vero cells (ATCC, Manassas, VA). RNA-sequencing was applied to confirm the full-length JURV genome (10,993 bp) as previously described.^4^ Infectious JURV was recovered from a full-length cDNA clone (Genscript, USA) comprising genes encoding for the nucleoprotein (JURV-N), phosphoprotein (JURV-P), matrix protein (JURV-M), glycoprotein (JURV-G), and RNA-directed RNA polymerase L protein (JURV-L) as described by Lawson et al.^14^ Vesicular stomatitis virus (VSV) was rescued from the pXN2 cDNA plasmid, and virus stock was amplified on BHK-21 cells. Sucrose density gradient centrifugation was used to obtain purified viral particles (VSV, MORV and recombinant JURV) before *in vitro* and *in vivo* studies.

### Amplification of viral stock

Viral amplification was done by infecting confluent (∼80%) Vero cells in T-175 flasks with a low multiplicity of infection (MOI) of 0.001 of JURV, MORV, or VSV. Forty-eight hours post-infection or when cytopathic effects (CPE) were observable. Supernatants of virus-infected cells were collected from the flasks. The viral stocks were purified using 10-40% sucrose-density gradient ultracentrifugation followed by dialysis. The titer (TCID_50_) of each virus was determined by the Spearman-Kärber algorithm using serial viral dilutions in BHK-21 cells.

### Cell viability assays

For all cytotoxicity assays (96-well format), 1.5 x 10^4^ cells were infected with JURV, MORV, or VSV at the indicated MOI of 10, 1, and 0.1 in serum-free Gibco Minimum Essential Media (Opti-MEM). Cell viability was determined using Cell Titer 96 AQueous One Solution Cell Proliferation Assay (Promega Corp, Madison, Wisconsin, USA). Data was generated as means of six replicates from two independent experiments +/- SEM.

### Crystal violet assays

Five hundred thousand cells were infected with oncolytic JURV in 6-well plates at an MOI of 0.1 for 1h. Supernatants of virus-infected cells were removed, and cells were washed with PBS and incubated at 37°C until analysis. At 72 hours after infection, cells were fixed with 5% glutaraldehyde and stained with 0.1% crystal violet to visualize cellular morphology and remaining adherence indicative of cell viability. Pictures of representative areas were taken.

### One-step viral growth kinetics

Two hundred thousand HCC cells were plated in each well of a 6-well plate in 2 mL of complete DMEM. After overnight rest, we infected the cells with JURV at an MOI of 0.1 for 1 hour. Supernatants of virus-infected cells were removed, and cells were washed with PBS, and fresh media was added. At timepoints 10, 24, 48, and 72 hours, the supernatant was collected and stored at −80C. Viral titers (TCID_50_) were determined with serial dilutions of the supernatant on Vero cells. Data was generated as means of two independent experiments +/- SEM.

### Interferon sensitivity assays

HCC cells were seeded in a 96-well plate at a density of 2.0 x 10^4^ cells/well and cultured overnight. Twenty-four hours post-infection, cells were pretreated with different concentrations of Universal type I IFN-α was added directly into the culture medium. After overnight incubation, fresh medium containing Universal Type I IFN-α (Catalog No. 11105-1; PBL Assay Science, USA) was added, and cells were infected with JURV at an MOI of 0.01. Cell viability was assessed using a Cell Titer 96 AQueous One Solution Cell Proliferation Assay (Promega Corp, Madison, Wisconsin, USA). Absorbance measurements at 490 nm were normalized to the maximum read per cell line, representing 100% viability. Data are shown from three independent experiments. For all cell viability experiments, absorbance was read using a Cytation 3 Plate Reader (BioTeK, Winooski, VT, USA). Data are expressed as means of triplicates from three independent experiments +/- SEM.

### Mice

Female C57BL6/J (Strain #:000664) and NOD.Cg-*Prkdc^scid^*/J (Strain #:001303) were purchased at 6-8 weeks of age from the Jackson Laboratory. All mice were housed at the Division of Laboratory Animal Medicine (DLAM) at University of Arkansas for Medical Sciences (UAMS). The DLAM has a full staff of veterinarians and veterinary technicians who supervised and assisted in animal care throughout the studies. All animal studies were conducted in accordance with and approved by the Institutional Animal Care and Use Committee at the University of Arkansas for Medical Sciences.

### Analysis of virus-induced adverse events in mice

Female C57BL/6J mice (N=6 mice/group) were intranasally (25 µL in each nostril) or intravenously (50 µL/mouse) administered with phosphate-buffered saline (PBS), moderately high dose (1 x 10^7^ TCID_50_), or high dose (1 x 10^8^ TCID_50_) of the virus. Body weight, temperature, behavior, and clinical signs were monitored by a board-certified veterinarian at least three times a week to detect any signs of toxicity. However, three days post-infection were sacrificed, three mice per group and tissues were collected (blood, brain, liver, and spleen) for short-term toxicity evaluation and viral biodistribution. The remaining mice were monitored for thirty days.

### Short-term toxicological analysis of blood components

Blood was collected from the submandibular vein (cheek bleed) and cardiac puncture on day 3 post-treatment. Blood was collected in BD Microtainer tubes with ethylenediaminetetraacetic acid or lithium heparin (Becton, Dickinson and Company, Franklin Lakes, New Jersey, USA) for complete blood counts (CBC) or in BD Microtainer SST tubes (Becton, Dickinson, and Company) for serum analysis. CBC analysis was performed in an Abaxis Piccolo Xpress chemistry analyzer (Abaxis, Union City, California, USA), and blood chemistry analysis was done in a VetScan HM5 Hematology Analyzer (Abaxis).

### Toxicoproteomic analysis

At three days post-inoculation of JURV, mouse brain and liver tissues were harvested and dehydrated using an increasing ethanol concentration and embedded into paraffin to become formalin-fixed paraffin-embedded (FFPE) blocks as previously described.^96^ Tissue blocks were sectioned into 3-5 10 µm sections and underwent a deparaffinization procedure for FFPE tissue.^97^ Following deparaffinization of FFPE samples with xylene and tissue lysis in sodium dodecyl sulfate, total protein was reduced, alkylated, and digested using filter-aided sample preparation ^98^ with sequencing grade modified porcine trypsin (Promega). Tryptic peptides were separated by reverse-phase XSelect CSH C18 2.5 µm resin (Waters) on an in-line 150 × 0.075 mm column using an UltiMate 3000 RSLCnano system (Thermo). Peptides were eluted using a 60 min gradient from 98:2 to 65:35 (buffer A, 0.1% formic acid, 0.5% acetonitrile: buffer B, 0.1% formic acid, 99.9% acetonitrile) ratio. Eluted peptides were ionized by electrospray (2.4 kV) followed by mass spectrometric (MS) analysis on an Orbitrap Exploris 480 mass spectrometer (Thermo). MS data were acquired using a Fourier transform MS (FTMS) analyzer in profile mode at a resolution of 120,000 over a range of 375 to 1500 m/z. Following HCD activation, MS/MS data were acquired using the FTMS analyzer in centroid mode at a resolution of 15,000 and normal mass range with normalized collision energy of 30%. Proteins were identified by database search using MaxQuant (Max Planck Institute) label-free quantification with a parent ion tolerance of 2.5 ppm and a fragment ion tolerance of 20 ppm. Scaffold Q+S (Proteome Software) was used to verify MS/MS-based peptide and protein identifications. Protein identifications were accepted if they could be established with less than 1% false discovery and contained at least two identified peptides. Protein probabilities were assigned by the Protein Prophet algorithm.^99^

### In vivo efficacy of the oncolytic JURV in a syngeneic mouse model of HCC

To evaluate the in vivo therapeutic efficacy of oncolytic JURV in a syngeneic mouse HCC model, 1 x 10^6^ HEPA 1-6 cells in 100 µL of cold RPMI were injected subcutaneously into the right flanks of immunocompetent female C57BL6/J mice (n=7/group; Jackson Laboratory) using 1 mL syringes. Mice were monitored weekly for palpable tumors or any changes in appearance or behavior. When average tumors reached a treatable size (80-120 mm^3^), mice were randomized into the respective study groups and dosed within 24 hours of randomization. On days 0, 7, and 14, mice were administered 50 µL IT injections containing PBS (control group) or 1 x 10^7^ TCID_50_ units of JURV (Test article group). Groups administered anti-PD-1 therapy or combination JURV + anti-PD-1 also receive 50 µL of anti-PD-1 antibodies intraperitoneally (IP) twice a week for three weeks. To establish the syngeneic bilateral (dual flanks) HCC tumors, HEPA 1-6 cells (1 x 10^6^ cells/mouse) were first subcutaneously grafted into the right flanks. These resulted in tumors in approximately 14 days and were categorized as “primary” tumors. Simultaneously, we performed distant HEPA 1-6 tumor grafts injection (1 x 10^6^ cells/mouse) into the left flanks of these mice. Mice in the dual flank group received 50 µL IT injections of 1 x 10^7^ TCID_50_ units of JURV on their right flanks only once a week for three weeks. Tumor volume and body weight were measured twice weekly following randomization and initiation of treatment using a digital caliper and balance. Tumor volume was calculated using the following equation: (longest diameter * shortest diameter2)/2 with a digital caliper. During the first week of treatment and after each injection, mice were monitored daily for signs of recovery for up to 72 hours. Mice were euthanized when body weight loss exceeded 20%, when tumor size was larger than 2,000 m³ or for adverse effects of treatment. Mice were sacrificed 28 days following the first JURV dose administration, at which time tumor and blood were collected for downstream analysis.

### HEP3B Xenograft Model

Female NOD.Cg-*Prkdc^scid^*/J mice were subcutaneously inoculated with HEP3B cells tagged with a firefly luciferase reporter gene on the right flanks (n=6/group). When the average tumor volume reached 80-120 mm^3^, mice were administered 50 µL IT injections of 1 x 10^7^ TCID_50_ of JURV or 50 mL of PBS weekly for three weeks. Tumor volume was measured twice weekly until the end of the study (Day 21), or the humane endpoint as described above. We also recorded mouse body weight and clinical observations twice per week.

### Bioluminescence imaging

Tumor-bearing (HEP3B) mice were anesthetized with isoflurane and imaged once a week (days 0, 7, and 14) with the IVIS Xenogen imaging system to virus-induced changes in tumor growth. Anesthesia was induced in an induction chamber (2–5% isoflurane), after which the mice were placed in the imaging instrument and fitted with a nose cone connected to a vaporizer to maintain isoflurane (1.5–2%) during the procedure. This range of concentrations produces a level of anesthesia that prevents animal movement during scanning. If the respiratory rate accelerates or slows, the isoflurane concentration is increased or decreased. We used a heated animal bed, heating pads, and, if necessary, a heating lamp to ensure that body temperature is maintained both before imaging and during the procedure. Each mouse received an intraperitoneal injection of D-luciferin (Sigma-Aldrich # L9504; 150 mg/kg body weight in the volume of 10 µl/g of body weight, prepared in sterile water). Anesthetized mice were placed into the IVIS Xenogen imaging system on their stomachs. Imaging of each group of mice took less than 10 minutes. This is a non-invasive imaging procedure, and we needed no restraints.

### Analysis of tumor-infiltrating immune cells

Hepa 1-6 tumors (n= 3 samples/group) were excised and dissociated using a mouse tumor dissociation kit (Miltenyi, CAT# 130-096-730) with a gentleMACS^™^ Octo Dissociator (Miltenyi) according to the manufacturer’s protocol. CD45^+^ cells were isolated with mouse CD45 (TIL) microbeads (Miltenyi). Cells were incubated with Fixable Viability Stain 510 for 15 minutes at *4 °C* followed by anti-Fc blocking reagent (Biolegend, Cat# 101320) for 10 minutes prior to surface staining. Cells were stained, followed by data acquisition with a BD LSRFortessa X-20 flow cytometer. All antibodies were used following the manufacturer’s recommendation. Fluorescence Minus One control was used for each independent experiment to establish gating. For intracellular staining of granzyme B, cells were stained using an intracellular staining kit (Miltenyi), and analysis was performed using FlowJo™ (TreeStar). Forward scatter and side scatter cytometry were used to exclude cell debris and doublets.

### Flow cytometry antibody analysis

The following antibodies were used for flow cytometry analysis: CD45-FITC (Cat. # 553079; *BD Biosciences), CD3-BUV395 (Cat. # 563565; BD Biosciences), CD4-BUV737 (Cat. # 612761; BD Biosciences), CD8-Percp-Cy5.5 (Cat. # 45-0081-82; eBioscience), CD44-BV711 (Cat. # 103057; Biolegend), CD335-PE/Dazzle594 (Cat. #137630; Biolegend), PD-1-PE (Cat. # 551892; BD Biosciences), Ki67*-BV605 (Cat. # 652413; Biolegend), Granzyme B*-APC (Cat. # 366408; Biolegend),* IFN-γ*-BV421 (Cat. # 563376; BD Biosciences), CD11b-PE-Cy7 (Cat. # 101216; Biolegend), F4/80-BV510 (Cat. # 123135; Biolegend), CD206-AF700 (Cat. # 141734; Biolegend), I-A/I-E-BV786 (Cat. # 743875; BD Biosciences), and L/D-efluor780 (Cat. # 65-0865-18; eBioscience).

### RNA-sequencing of murine HCC tumors

Hepa 1-6 (n=3 samples/group) FFPE scrolls were processed for DNA and RNA extraction using a Quick-DNA/RNA FFPE Miniprep Kit with on-column DNase digestion for the RNA preps (Cat. # R1009; Zymo Research). RNA was assessed for mass concentration using the Qubit RNA Broad Range Assay Kit (Cat. # Q10211; Invitrogen) with a Qubit 4 fluorometer (Cat. # Q33238; Invitrogen). RNA quality was assessed with a Standard Sensitivity RNA Analysis Kit (Cat. # DNF-471-0500; Agilent) on a Fragment Analyzer System (Cat. # M5310AA; Agilent). Sequencing libraries were prepared using TruSeq Stranded Total RNA Library Prep Gold (Cat. # 20020599; Illumina). RNA DV200 scores were used to determine fragmentation times. Libraries were assessed for mass concentration using a Qubit 1X dsDNA HS Assay Kit (Cat. # Q33231; Invitrogen) with a Qubit 4 fluorometer (Cat. # Q33238; Invitrogen). Library fragment size was assessed with a High Sensitivity NGS Fragment Analysis Kit (Cat. # DNF-474-0500; Agilent) on a Fragment Analyzer System (Cat. # M5310AA; Agilent). Libraries were functionally validated with a KAPA Universal Library Quantification Kit (Cat. # 07960140001; Roche). Sequencing was performed to generate paired-end reads (2 × 100 bp) with a 200-cycle S1 flow cell on a NovaSeq 6000 sequencing system (Illumina).

### Bioinformatics analysis

We examined the mRNA and protein expression profiles of Hepa 1-6 tumors treated with PBS, JURV, anti-PD-1 or JURV + anti-PD-1. Three replicates were used to analyze each of the untreated (PBS) and treatment groups. The tumor samples were sequenced on an NGS platform. The files containing the sequencing reads (FASTQ) were then tested for quality control (QC) using MultiQC.^100^ The Cutadapt tool trims the Illumina adapter and low-quality bases at the end. After the quality control, the reads were aligned to a mouse reference genome (mm10/GRCm38) with the HISAT2 aligner^101^, followed by counting reads mapped to RefSeq genes with feature counts. We generated the count matrix from the sequence reads using HTSeq-count.^102^ Genes with low counts across the samples affect the false discovery rate, thus reducing the power to detect differentially expressed genes; thus, before identifying differentially expressed genes, we filtered out genes with low expression utilizing a module in the limma-voom tool^32^. Then, we normalized the counts by using TMM normalization^103^, a weighted trimmed mean of the log expression proportions used to scale the counts of the samples. Finally, we fitted a linear model in limma to determine differentially expressed genes and expressed data as mean ± standard error of the mean. All *p* values were corrected for multiple comparisons using Benjamini-Hochberg FDR adjustment. After identifying differentially expressed genes, enriched pathways were performed using the Ingenuity Pathway Analyses tool to gain biological insights. The statistical difference between groups was assessed using the nonparametric Mann-Whitney U test R module.

### Integration of transcriptomics and proteomics

The limma-normalized transcript expression levels and the normalized protein intensities were integrated using two independent methods. Firstly, the mixOmics package (Omics Data Integration Project R package, version 6.1.1) was implemented to generate heatmaps of the associated DEPs/DEGs as previously described.^104^ Secondly, the MOGSA package was used to generate heatmaps of the top 30 up- or down-regulated DEPs/DEGs between the various groups.^105^

### Blood chemistry and cytokines

Blood chemistry analysis was performed with an Abaxis Piccolo Xpress chemistry analyzer (Abaxis) to assess liver toxicity (i.e., aspartate transaminase, alkaline phosphatase, albumin), nephrotoxicity (i.e., creatinine, blood urea nitrogen), and serum electrolytes. Murine type I interferon-beta assay was performed using Mouse IFN beta SimpleStep ELISA® Kit (Cat. # ab252363; Abcam).

### TUNEL assay immunohistochemistry

Tumor tissue sections were subjected to the terminal deoxynucleotidyl transferase deoxyuridine triphosphate nick-end labeling (TUNEL) assay using the In Situ Cell Death Detection Kit (Roche Diagnostics, Indianapolis, IN) according to the manufacturer’s protocol. After staining, cells were counterstained with 4’,6-diamidino-2-phenylindol (DAPI) to visualize cell nuclei, mounted under cover slips with Prolong® Antifade kit (Invitrogen, Carlsbad, CA) and acquired using the Olympus IX-81 inverted microscope (Olympus America, Center Valley, PA) equipped with Hamamatsu ORCA-ER monochrome camera (Hamamatsu Photonics K.K., Hamamatsu City, Japan). Image analysis was performed using SlideBook 6.2 software. For quantification, 10 independent fields of view were collected per each well (each n) and mean optical density (MOD) or area of colocalization in pixels were recorded for Fluorescein (TUNEL) channel.

### Statistical analysis

All values were expressed as the mean ± standard error of mean, and the results were analyzed by one-way analysis of variance followed by the Tukey test or Benjamini-Hochberg FDR adjustment for multiple comparisons and *t* test to compare group means. Kaplan-Meier method for survival, using statistical software in GraphPad Prism, version 8 (GraphPad Software). A *p* value less than 0.05 was considered statistically significant.

### Data deposition

The RNA sequencing data are freely available via GEOGSE199131, and the proteomics data are available via ProteomeXchange with identifier PXD035806.

## Acknowledgments

We thank the personnel of the DNA Damage and Toxicology, Proteomics, Genomics, and Bioinformatic Cores at the University of Arkansas for Medical Sciences (UAMS) for their assistance during these studies.

## Author contributions

YZ, AB, MJC and BMN contributed to study concept and design, data acquisition, data analysis, data interpretation, and manuscript drafting. MJB, MT, ALG, KUF, MG, MT, CSS, JCC, CD, OB, SRP, and TK contributed to drafting and critical revision of the manuscript. CLW, DA, AG, SDB contributed to bioinformatic analysis. All authors approved the final, submitted version of the manuscript.

## SUPPLEMENTAL FIGURES AND TABLES

**Supplemental Figure 1.** S*c*hematic *representation of the antigenomic structure of JURV, VSV, and MORV.* The antigenome of JURV is typical of that of Rhabdovirus. Schematic representation of the 3’ to 5’ antigenomic organization of **a**) attenuated jurona virus (JURV), **b**) Wild-type vesicular stomatitis virus (VSV), and **c**) Morreton virus (MORV).

**Supplemental Figure 2.** Toxicologic pathology examination and dose-dependent response to treatment with type interferon on the oncolytic activity of JURV. Effect of exogenous IFN-β on the outcome of JURV-induced tumor cell death. Infectious titers and MOI dependent responses to species-specific exogenous type I IFN-β on HEP3B and HEPA 1-6 cell lines used as tumor models (**a**) HEP3B and (**b**) HEPA 1-6 cell lines. Results are from three independent experiments and are plotted as mean ± SEM. Intranasal (IN, brain) and intravenous (IV, liver and spleen) administration of 1 x 10^7^ or 1 x 10^8^ TCID_50_ of JURV in non-tumor bearing mice. (**c**) Hematoxylin and eosin (H&E) stained brain, liver and spleen sections from mice treated with low (1 x 10^7^ TCID_50_) or high dose (1 x 10^8^ TCID_50_) of oncolytic JURV.

**Supplemental Figure 3.** Toxicoproteomic analysis of differentially expressed proteins in brain and liver tissues of mice injected with low doses of oncolytic JURV in healthy non-tumor bearing. Volcano plot of protein expression differences for the brain (**a**) and liver (**c**) of mice treated with PBS vs. 1 x10^7^ TCID_50_ of JURV. 3-D Pie slices of the numbers of differentially expressed proteins (DEPs) for the brain (**b**) and liver (**d**) of mice injected with PBS vs. 1 x10^7^ TCID_50_ of JURV. Heatmaps of the top 20 DEPs up or down regulated in the brain (**c**) and liver (**d**) of mice injected with PBS vs. 1 x10^7^ TCID_50_ of JURV. DEPs were determined using the limma-voom method. A fold-change |*log*FC| ≥ 1 and a false discovery rate (FDR) of 0.055 were used as a cutoff. The *log*FC was computed using the difference between the mean of *log2*(JURV) and the mean of *log2*(PBS), that is, mean of *log2*(JURV) -mean of *log2*(PBS). Graph showing top-scoring canonical pathways significantly enriched by treatment with 1 x10^7^ TCID_50_ of JURV in the brain (**e**) and liver (**f**).

**Supplemental Figure 4.** C*h*anges *in body weight and Survival of Hepa 1-6 tumor-bearing mice Treated with oncolytic JURV.* Individual tumor volumes of HEPA 1-6 tumor-bearing mice between various groups in the (**a**) combination therapy study and (**b**) abscopal effect study. (**c**) Body weight of HEPA 1-6 tumor-bearing mice between various groups in the combination therapy study and abscopal effect study. Kaplan–Meier survival analysis of HEPA 1-6 tumor-bearing mice in the (**d**) combination therapy study (**e**) and abscopal effect study. The statistical significance of differences in the survival curves between the groups was evaluated using the log-rank (Mantel-Cox) test.

**Supplemental Figure 5.** A*n*alysis *of biomarkers of drug-induced hepatotoxicity and nephrotoxicity in mice treated with oncolytic JURV.* Changes in the serum biomarkers of drug-induced toxicity, including albumin (**a**, ALB), alkaline phosphatase (**b**, ALP), alanine aminotransferase (**c**, ALT), amylase (**d**, AMY), total bilirubin (**e**,TILB), blood urea nitrogen (**f**, BUN), calcium (**g**, CA), phosphate (**h**, PHOS), creatinine (**i**), glutamine (**j**, GLU), sodium (**k**, NA+), potassium (**l**, K+), total protein (**m**, TP), serum globulin (**n**, GLOB) and serum hemoglobulin (**o**, HEM).

**Supplemental Figure 6.** A*n*alysis *of tumor-infiltrating immune cells following intratumoral injection of oncolytic JURV in murine HCC tumors.* Immune profiling of PBS, JURV, anti-PD-1 therapy, JURV + anti-PD-1 antibodies and dual flanks (abscopal effect) treated syngeneic HEPA 1-6 mice showing percentage of tumor infiltrating immune cells, including natural killer T (**a**, NKT), F4/80-macrophages (**b**), CD45 + (**c**), CD11b+ (**d**), CD11b-(**e**), CD3+ (**f**), CD8+ (**g**), CD8+ Granzyme+ (**h**), CD8+ IFNg+ (**i**), CD4+ CD44+ (**j**), CD4+ IFNg+ (**k**) and CD4+ KI67+ (**l**) cells. The Bartlett test was used to test homogeneity of variance and normality. If the *p* value of the Bartlett test was no less than *P*=0.05, ANOVA and a two-sample *t* test were used to compare group means. If the *P* value of the Bartlett test was less than 0.05, the Kruskal-Wallis and Wilcoxon rank-sum tests were used to compare group means. The figures demonstrate the potential significant difference of the gated subsets in the CD45+ population, determined by the *p* value.

**Supplemental Figure 7.** Murine type I IFN expression in HEPA 1-6 tumor-bearing mice injected with PBS, JURV, anti-PD-1 antibodies, and JURV + Anti-PD-1 antibodies. (**a**) Level of antiviral cytokine (IFN-β) was measured in serum from mice treated with PBS, JURV, anti-PD-1 therapy, JURV + anti-PD-1 antibodies and dual flanks.

**Supplemental Figure 8.** P*r*oteogenomic *changes in murine HCC injected with oncolytic JURV.* (**a**) Volcano plot of mRNA expression differences for PBS vs. JURV (1 x 10^7^ TCID_50_). (**b**) 3-D Pie slices of the numbers of differentially expressed genes (DEGs) between PBS vs. JURV. (**c**) Heatmap of the top 20 DEGs up or down regulated PBS vs. JURV. DEGs were determined using the limma-voom. (**d**) Graph showing top-scoring canonical pathways significantly enriched by treatment with PBS vs. JURV. A MixOmics supervised analysis was carried out between DEPs and DEGs based on Log2 fold change values. Log2 fold change of DEG × Log2 fold change of DEP > 0 with a *P-value* of DEG and DEP <0.05 were considered associated DEGs/DEPs. (**e**) DEG/DEP expression heatmap of the 30 most up-regulated and down-regulated features DEG/DEP in PBS vs. JURV.

**Supplemental Figure 9.** T*r*anscriptome *and proteome analysis of anti-PD-1 therapy in murine HCC tumors.* (**a**) Volcano plot of mRNA expression differences for PBS vs. anti-PD-1 therapy (10 mg/kg, twice a week for 3 weeks). (**b**) 3-D Pie slices of the numbers of differentially expressed genes (DEGs) between PBS vs. anti-PD-1 antibodies. (**c**) Heatmap of the top 20 DEGs up or down regulated PBS vs. anti-PD-1 antibodies. DEGs were determined using the limma-voom. (**d**) Graph showing top-scoring canonical pathways significantly enriched by treatment with PBS vs. anti-PD-1 antibodies. A MixOmics supervised analysis was carried out between DEPs and DEGs based on Log2 fold change values. Log2 fold change of DEG × Log2 fold change of DEP > 0 with a *P-value* of DEG and DEP <0.05 were considered associated DEGs/DEPs. (**e**) DEG/DEP expression heatmap of the 30 most up-regulated and down-regulated features DEG/DEP in PBS vs. anti-PD-1 antibodies.

**Supplemental Figure 10.** C*o*mmon *DEGs between PBS vs. JURV vs. PBS vs. Anti-PD-1 vs. PBS vs. JURV + Anti-PD-1 in murine HCC.* (a) Venn diagram showing the 35 DEGs in HEPA 1-6 tumors from the three data sets (PBS vs. Anti-PD-1, PBS vs. JURV + Anti-PD-1, PBS vs. JURV). (**b**) List of common DEGs (35) between the various groups.

**Supplemental Figure 11.** Top canonical pathways enriched by common DEGs between PBS vs. JURV vs. PBS vs. Anti-PD-1 vs. PBS vs. JURV + Anti-PD-1 in murine HCC. Top immune-related canonical pathways enriched by common DEGs (35) *from* the three data sets (PBS vs. Anti-PD-1, PBS vs. JURV + Anti-PD-1, PBS vs. JURV). Analysis was performed using ReactomeGSA (https://reactome.org/dev/analysis).

**Supplemental Figure 12.** T*o*p *canonical pathways enriched in JURV vs. Dual flanks (abscopal effect).* (**a**) Top immune-related canonical pathways enriched in comparison JURV vs. Dual flanks.

**Supplemental Figure 13.** I*n*dividual *HEP3B tumor volume and survival.* Individual tumor volume after treatment. Female NOD.Cg-Prkdcscid/J (Strain #:001303) mice (n=6/group) mice were inoculated subcutaneously with HEP3B cells tagged with a luciferase reporter protein. When tumor volume was between 80-120 mm^3^, mice were divided into two groups and received IT injections with either PBS or JURV at a dose of 1 x 10^7^ TCID_50_ (Days 0,7 and 14). (**a**) Tumor volume and (**b**) body weight was recorded twice weekly until humane endpoint or end of study (Day 21). (**c**) Kaplan–Meier survival analysis of HEP3B tumor-bearing mice. The statistical significance of differences in the survival curves between the groups was evaluated using the log-rank (Mantel-Cox) test.

