## Supplemental Figures for "In vivo Safety and Immunoactivity of Oncolytic Jurona Virus in Hepatocellular Carcinoma: A Comprehensive Proteogenomic Analysis"

**Supplemental Figure 1.** Schematic representation of the antigenomic structure of JURV, VSV, and MORV

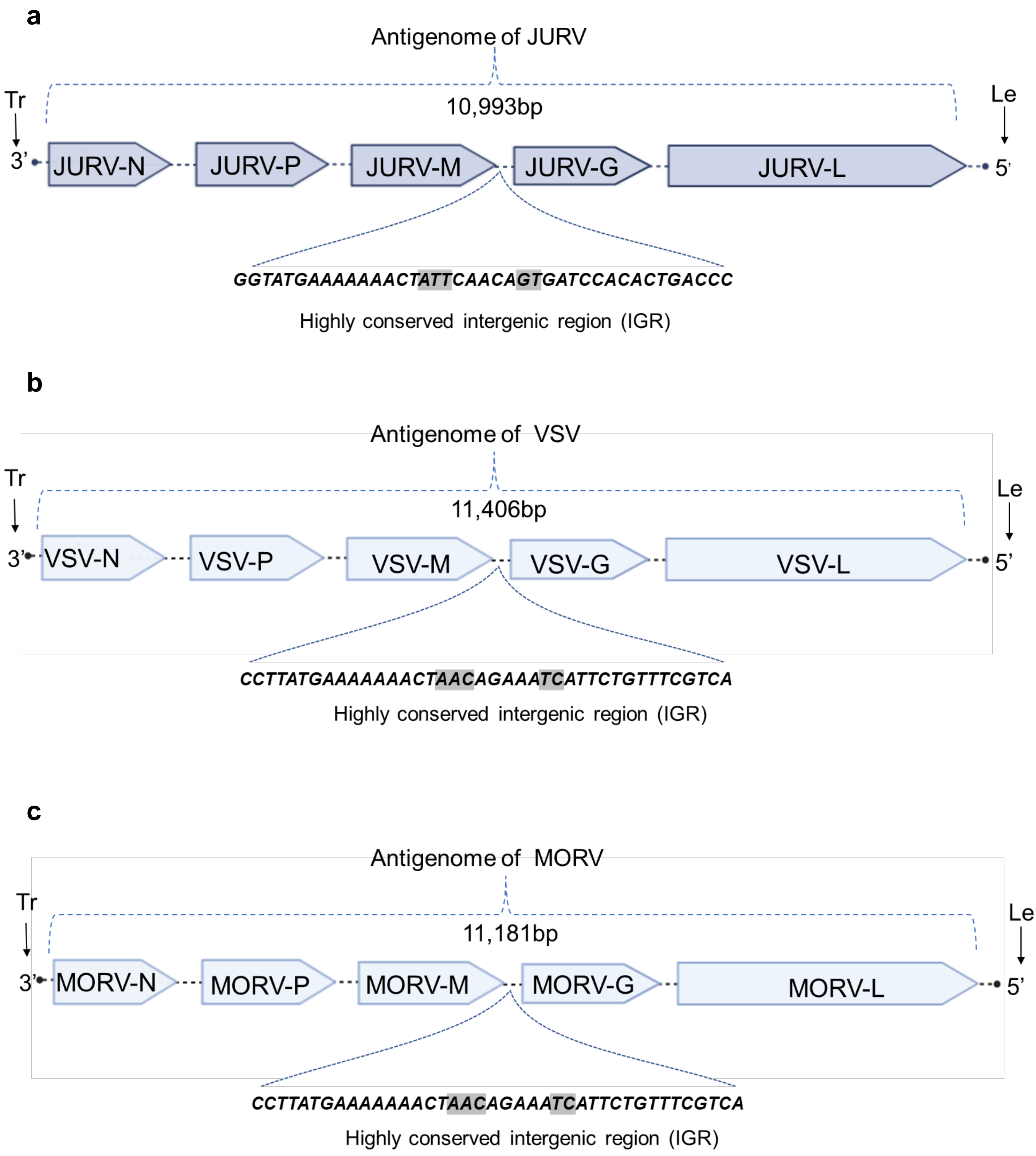

**Supplemental Figure 2**. Toxicologic pathology and toxicoproteomic examination of brain, liver, and spleen from mice injected (IV vs. IN) with low and high doses of oncolytic JURV

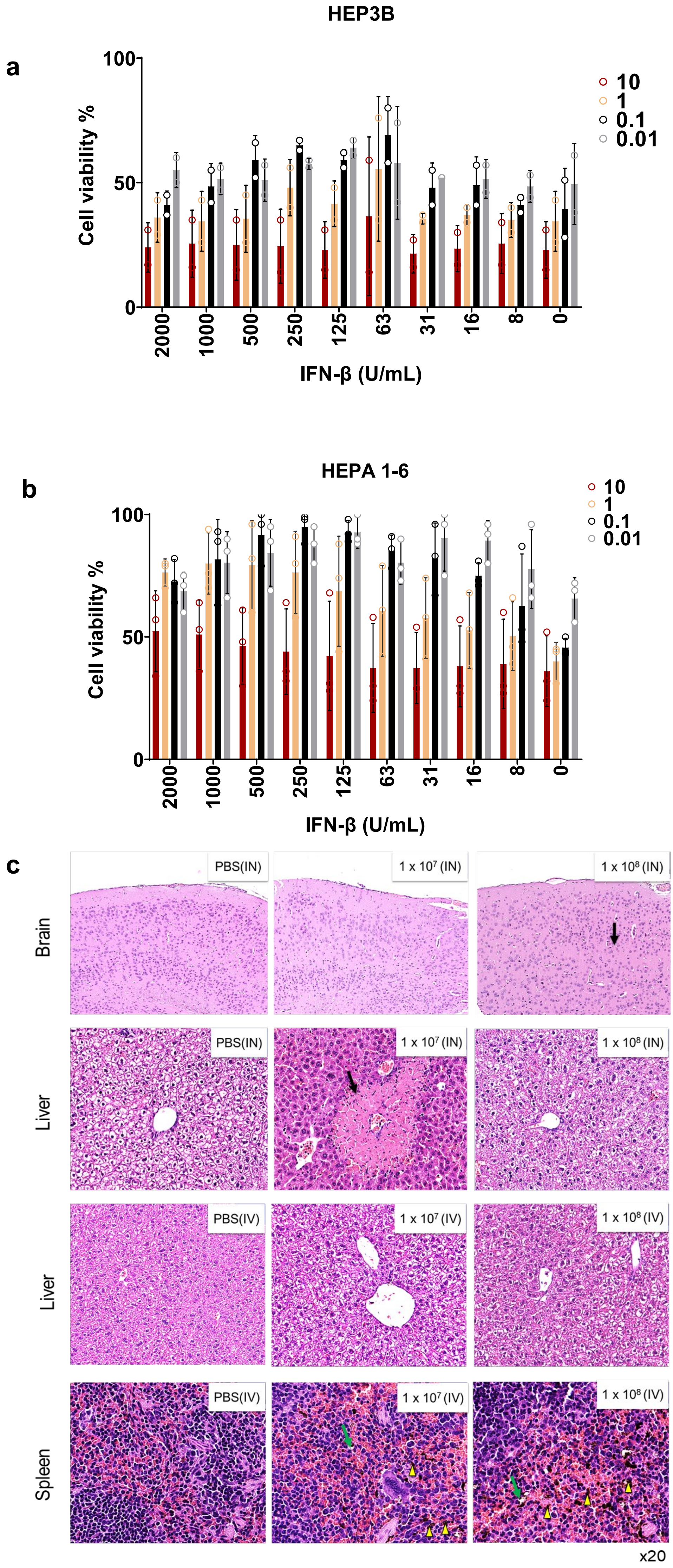


Black arrows: Indicate that samples are were within normal limits. Green arrows: Indicated necrosis, single cell, macrophage, sporadic. Yellow triangle: Indicated pigmentation increased in macrophages, red pulp and white pulp. (IN: Intranasal: IN: Intravenous)

**Supplemental Figure 3**. Toxicoproteomic analysis of differentially expressed proteins in brain and liver tissues of mice injected with low doses of oncolytic JURV in healthy non-tumor bearing mice

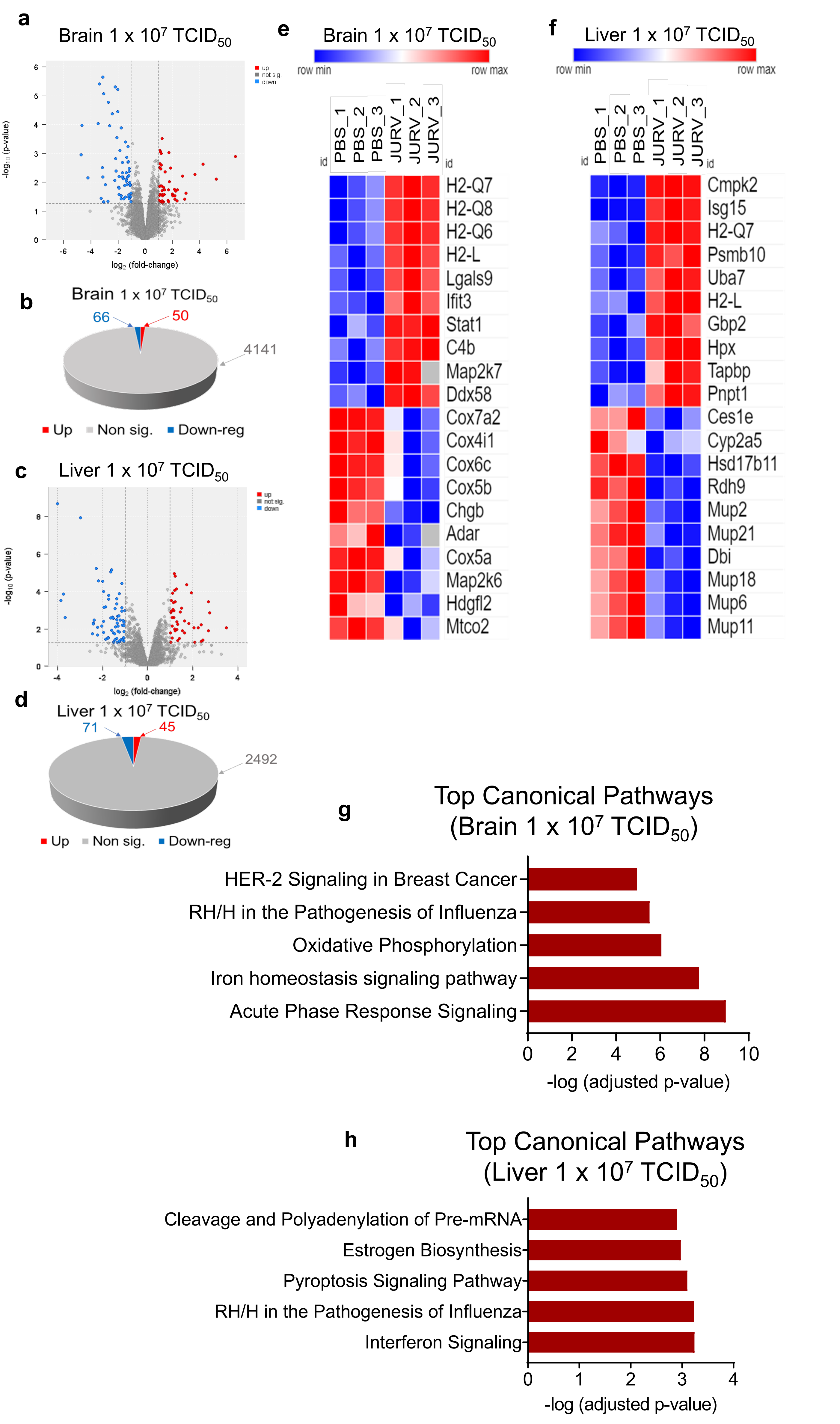

**Supplemental Figure 4**. Changes in body weight and Survival of Hepa 1-6 tumor-bearing mice Treated with oncolytic JURV

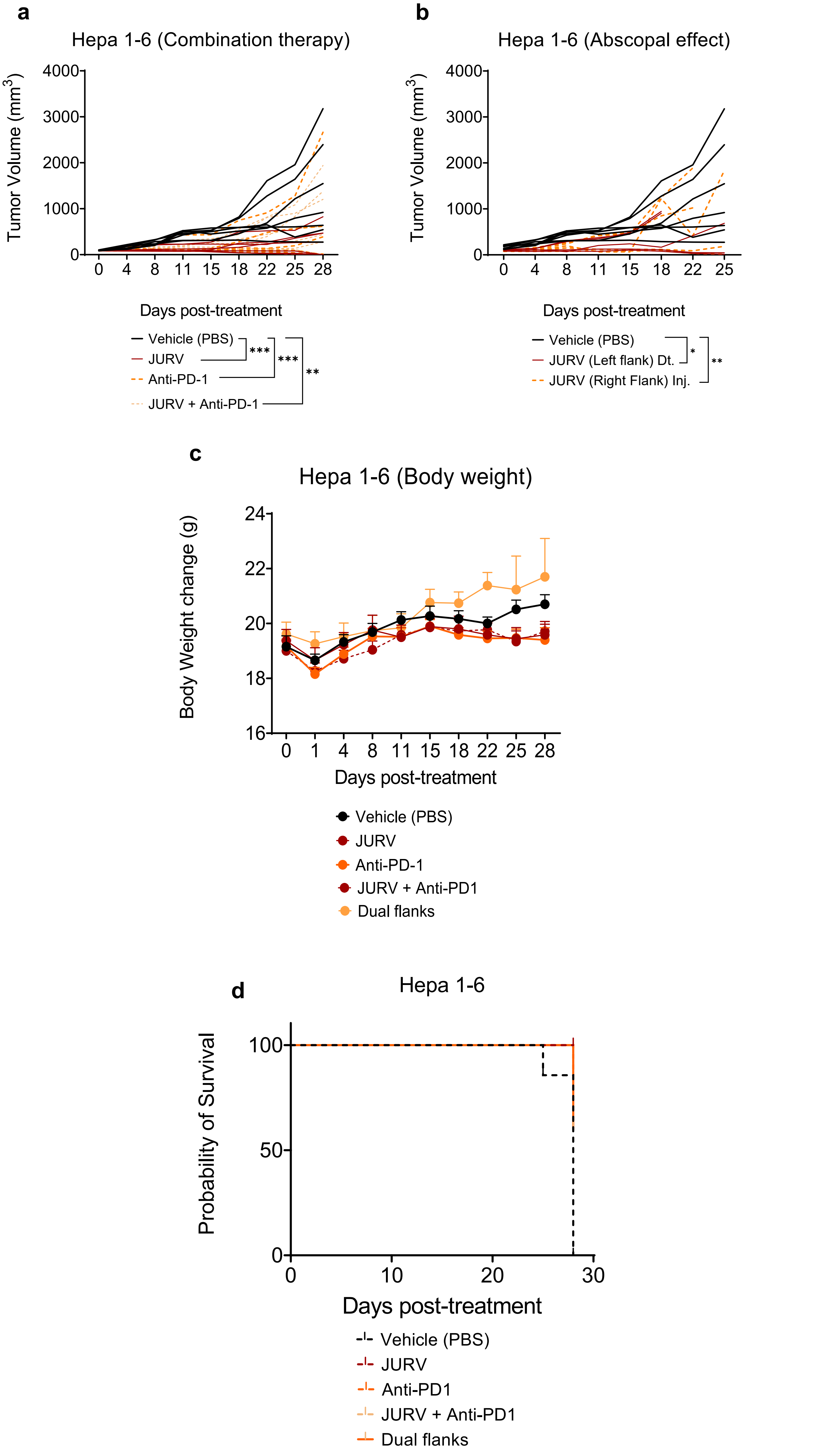

**Supplemental Figure 5**. Analysis of biomarkers of drug-induced hepatotoxicity and nephrotoxicity in mice treated with oncolytic JURV

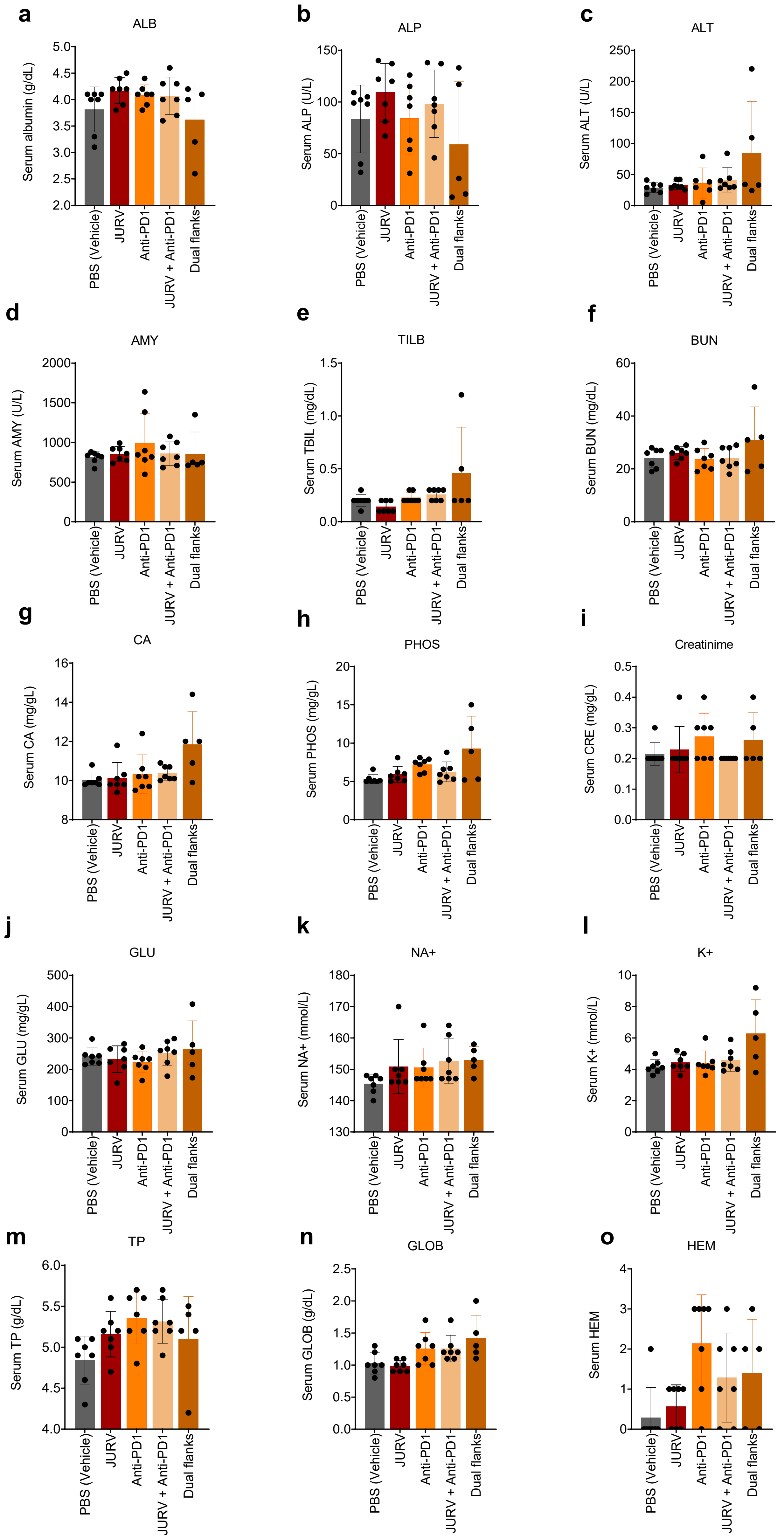

**Supplemental Figure 6**. Analysis of tumor-infiltrating immune cells following intratumoral injection of oncolytic JURV in murine HCC tumors

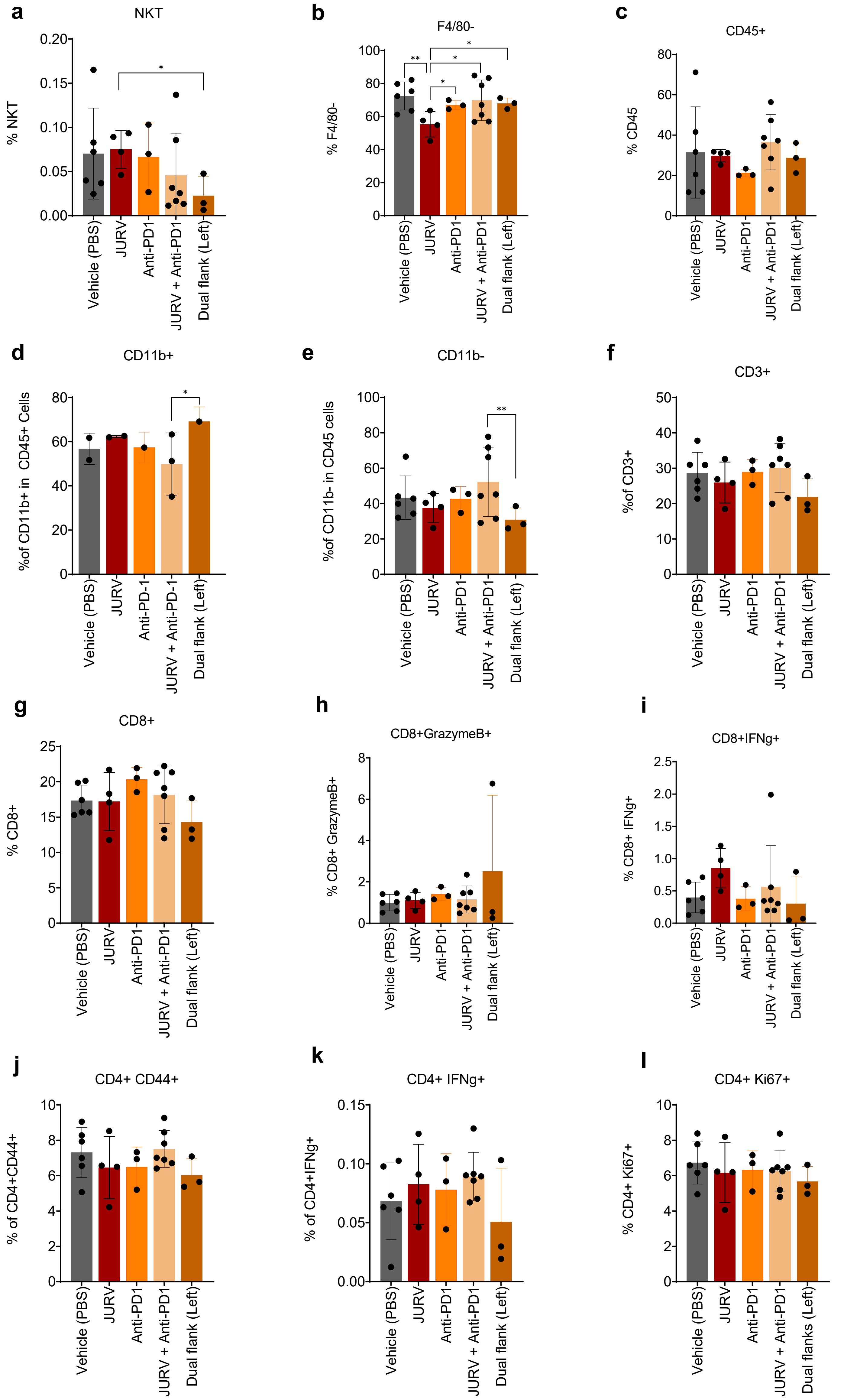

**Supplemental Figure 7.** Murine type I IFN expression in HEPA 1-6 tumor-bearing mice injected with PBS, JURV, anti-PD-1 antibodies, and JURV + Anti-PD-1 antibodies

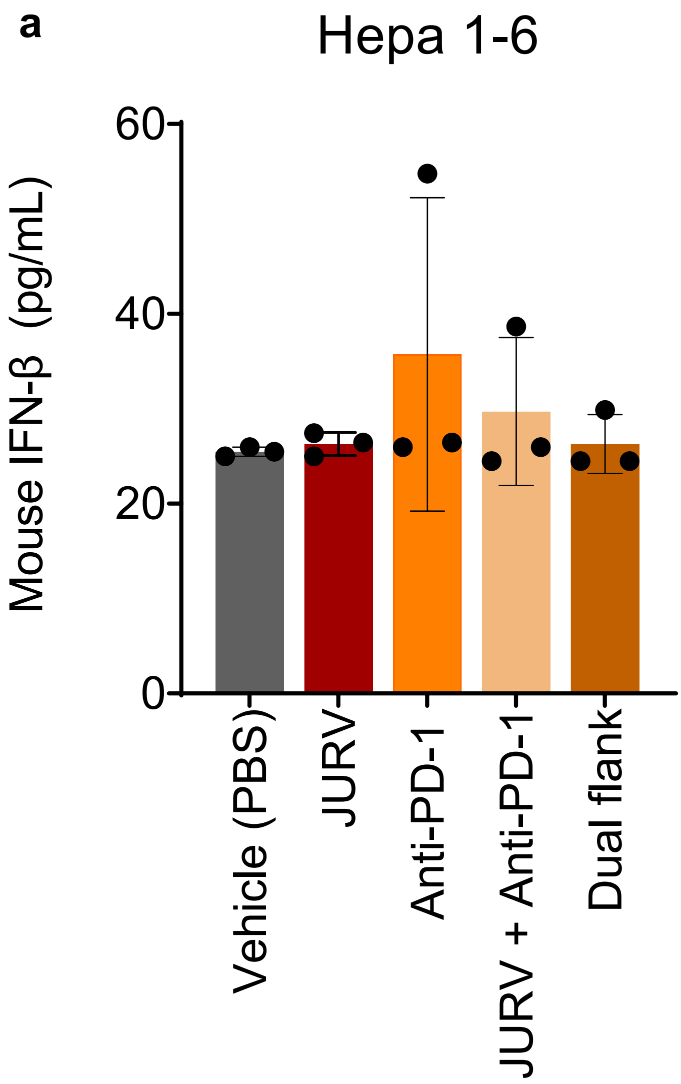

**Supplemental Figure 8**. Proteogenomic changes in murine HCC injected with oncolytic JURV

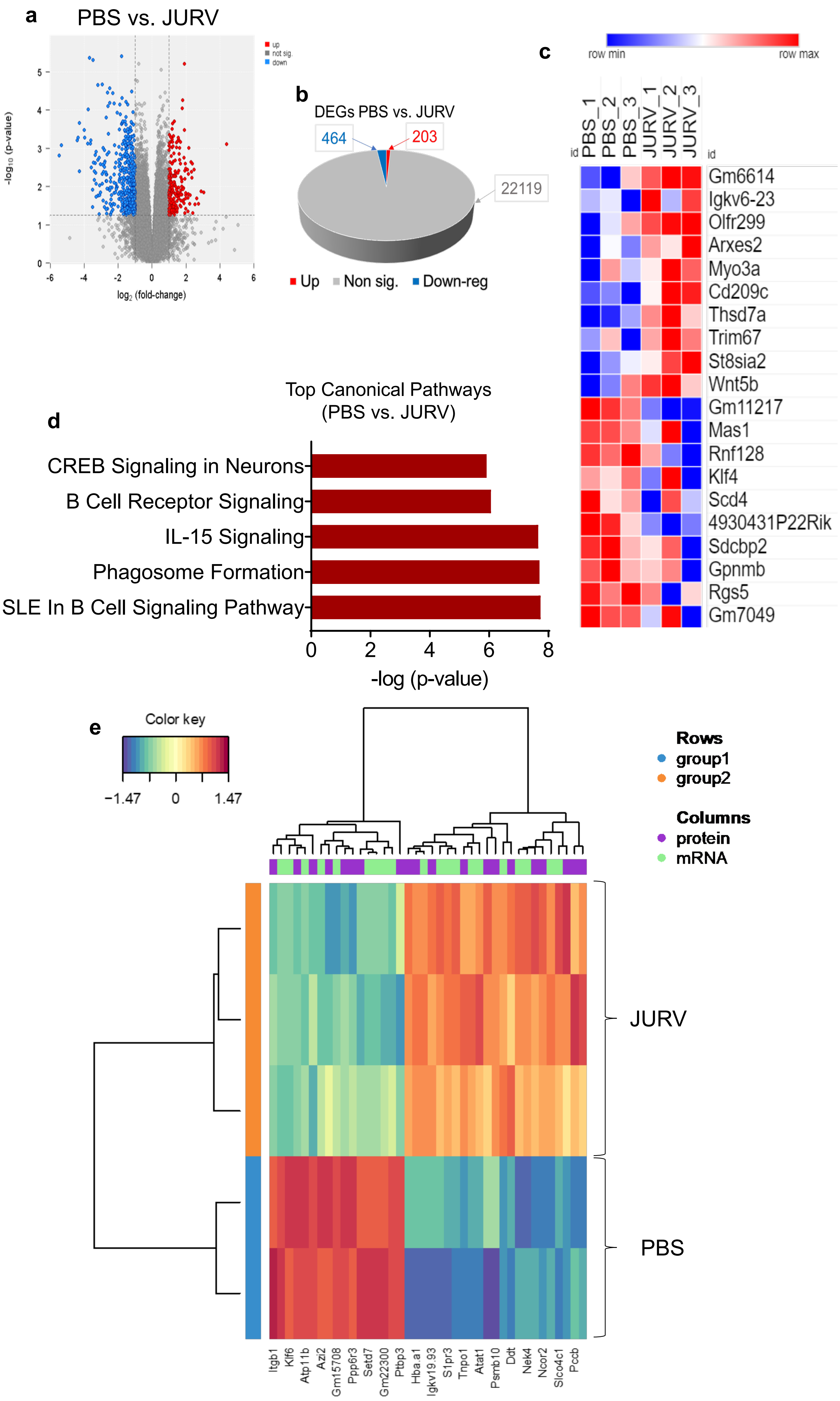

**Supplemental Figure 9**. Transcriptome and proteome analysis of anti-PD-1 therapy in murine HCC tumors

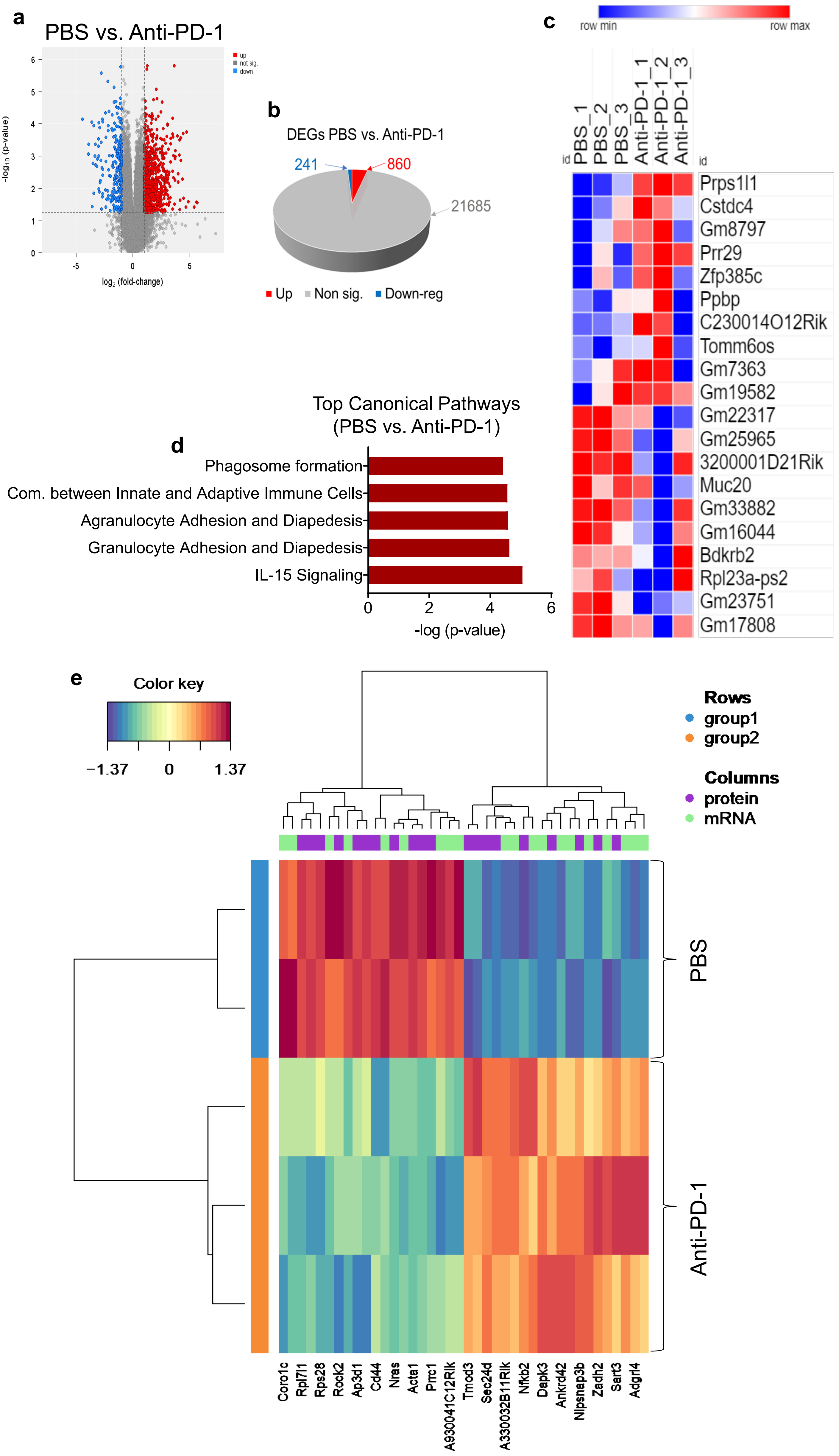

**Supplemental Figure 10.** Common DEGs between PBS vs. JURV vs. PBS vs. Anti-PD-1 vs. PBS vs. JURV + Anti-PD-1 in murine HCC

**a**

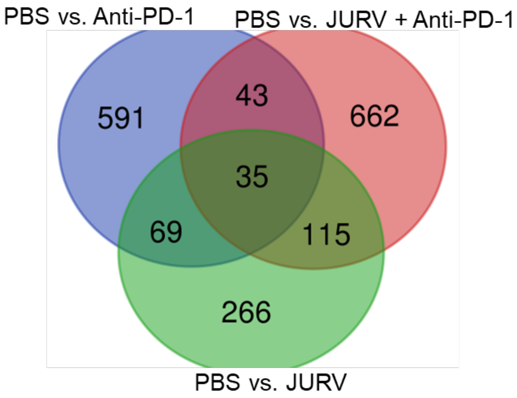


**b**

| Symbol | Entrez Gene Name | GenBank/ Gene Symbol | Location |
|---|---|---|---|
| ABCA12 | ATP binding cassette subfamily A member 12 | Abca12 | Plasma Membrane |
| AI463229 | expressed sequence AI463229 | AI463229 | Other |
| BCAS1 | brain enriched myelin associated protein 1 | Bcas1 | Plasma Membrane |
| CLVS1 | clavesin 1 | Clvs1 | Cytoplasm |
| CNTN5 | contactin 5 | Cntn5 | Plasma Membrane |
| COLEC11 | collectin subfamily member 11 | Colec11 | Extracellular Space |
| CPS1 | carbamoyl-phosphate synthase 1 | Cps1 | Cytoplasm |
| CSTA | cystatin A | Cstdc4 | Cytoplasm |
| DNAH12 | dynein axonemal heavy chain 12 | Dnah12 | Cytoplasm |
| GDF1 | growth differentiation factor 1 | Gdf1 | Extracellular Space |
| Gm16202 | sorting nexin 4 pseudogene | Gm16202 | Other |
| Gm19582 | predicted gene, 19582 | Gm19582 | Other |
| Gm22317 | -- | Gm22317 | Other |
| Gm38403 | predicted gene, 38403 | Gm38403 | Other |
| Gm7001 | CDK5 regulatory subunit associated protein 3 pseudogene | Gm7001 | Other |
| INSM1 | INSM transcriptional repressor 1 | Insm1 | Nucleus |
| LRRN4 | leucine rich repeat neuronal 4 | Lrrn4 | Plasma Membrane |
| LRRTM4 | leucine rich repeat transmembrane neuronal 4 | Lrrtm4 | Extracellular Space |
| MACROD2 | mono-ADP ribosylhydrolase 2 | Macrod2 | Nucleus |
| mir-679 | microRNA 679 | Mir679 | Cytoplasm |
| MYO3A | myosin IIIA | Myo3a | Cytoplasm |
| NALCN | sodium leak channel, non-selective | Nalcn | Plasma Membrane |
| NTM | neurotrimin | Ntm | Plasma Membrane |
| PABPC1L | poly(A) binding protein cytoplasmic 1 like | Pabpc1l | Cytoplasm |
| PLA2G2F | phospholipase A2 group IIF | Pla2g2f | Extracellular Space |
| PRELID3A | PRELI domain containing 3A | Prelid3a | Cytoplasm |
| ROBO2 | roundabout guidance receptor 2 | Robo2 | Plasma Membrane |
| Slc25a2 | solute carrier family 25 (mitochondrial carrier, ornithine transporter) member 2 | Slc25a2 | Cytoplasm |
| SMAD9 | SMAD family member 9 | Smad9 | Nucleus |
| TMPRSS3 | transmembrane serine protease 3 | Tmprss3 | Plasma Membrane |
| TOGARAM2 | TOG array regulator of axonemal microtubules 2 | Togaram2 | Other |
| Trbv3 | T cell receptor beta, variable 3 | Trbv3 | Other |
| TRIM67 | tripartite motif containing 67 | Trim67 | Cytoplasm |
| TTLL11 | tubulin tyrosine ligase like 11 | Ttll11 | Cytoplasm |
| UCP3 | uncoupling protein 3 | Ucp3 | Cytoplasm |

**Supplemental Figure 11.** Top canonical pathways enriched by common DEGs between PBS vs. JURV vs. PBS vs. Anti-PD-1 vs. PBS vs. JURV + Anti-PD-1 in murine HCC

**a**

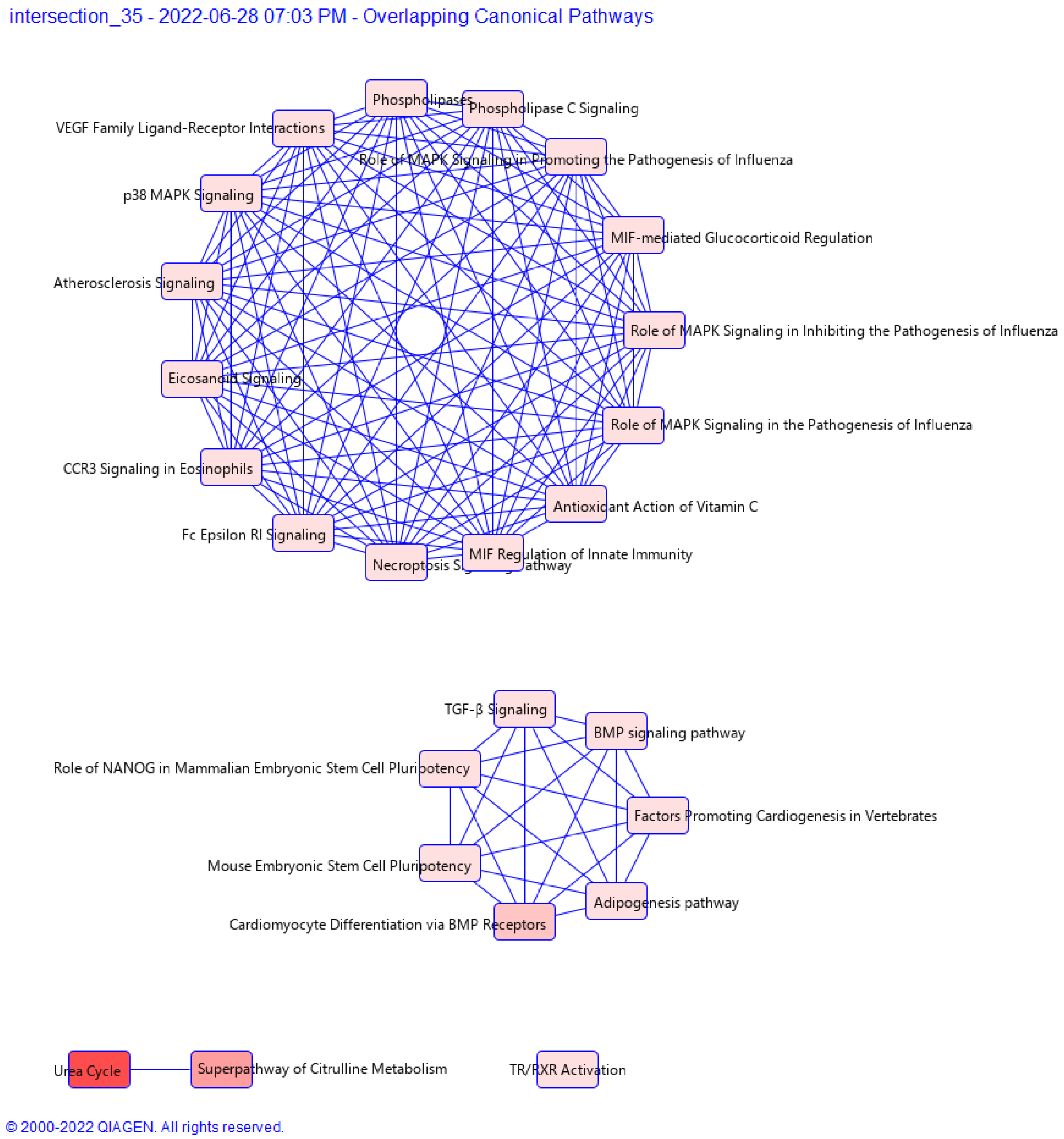

**Supplemental Figure 12.** Top canonical pathways enriched in JURV vs. Dual flanks (abscopal effect)

**a**

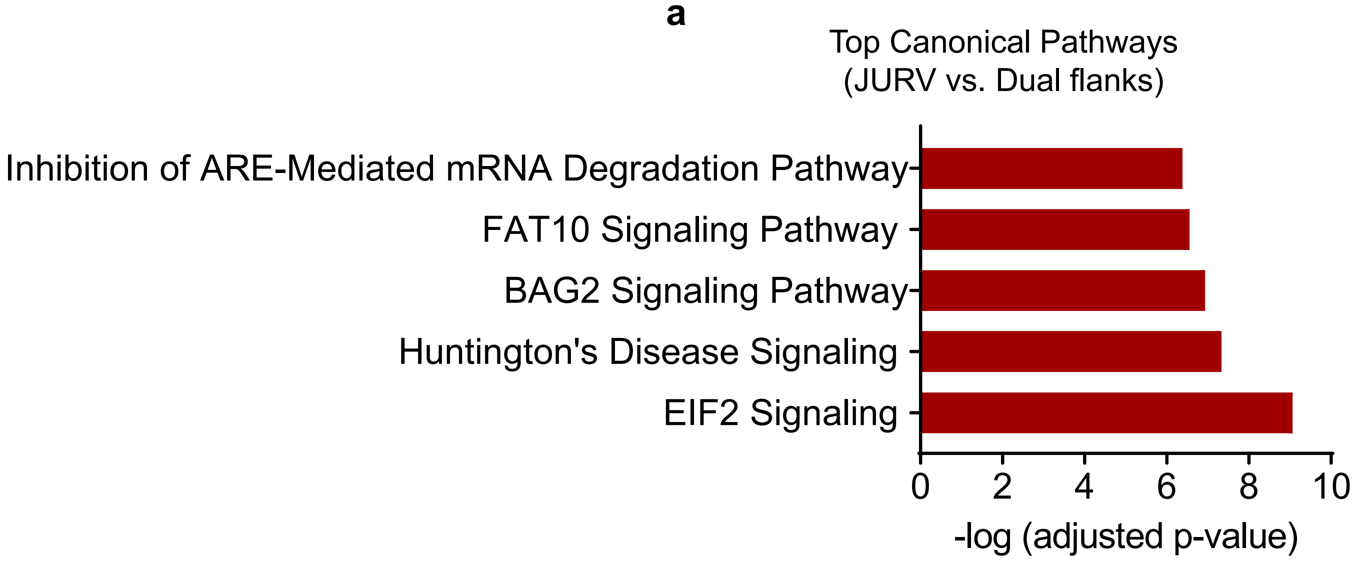

**Supplemental Figure 13**. Individual HEP3B tumor volume and survival

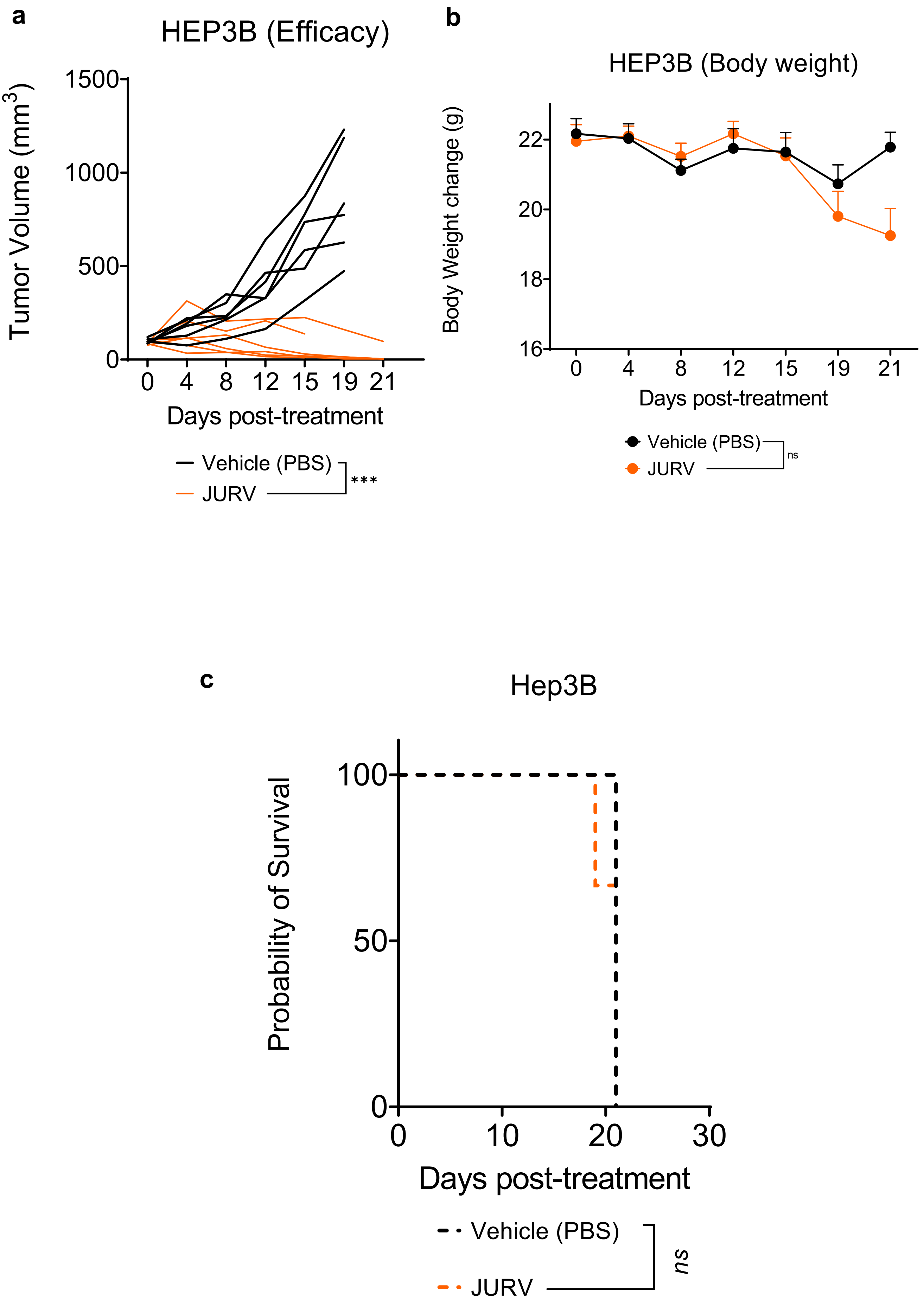
